# Increased yields of duplex sequencing data by a series of quality control tools

**DOI:** 10.1101/864835

**Authors:** Gundula Povysil, Monika Heinzl, Renato Salazar, Nicholas Stoler, Anton Nekrutenko, Irene Tiemann-Boege

## Abstract

Duplex sequencing is currently the most reliable method to identify ultra-low frequency DNA variants by grouping sequence reads derived from the same DNA molecule into families with information on the forward and reverse strand. However, only a small proportion of reads are assembled into duplex consensus sequences, and reads with potentially valuable information are discarded at different steps of the bioinformatics pipeline, especially reads without a family. We developed a bioinformatics tool-set that analyses the tag and family composition with the purpose to understand data loss and implement modifications to maximize the data output for the variant calling. Specifically, our tools show that tags contain PCR and sequencing errors that contribute to data loss and lower DCS yields. Our tools also identified chimeras, which result in unpaired families that do not form DCS. Finally, we also developed a tool called Variant Analyzer that re-examines variant calls from raw reads and provides different summary data that categorizes the confidence level of a variant call by a tier-based system. We demonstrate that this tool identified false positive variants tagged by the tier-based classification. Furthermore, with this tool we can include reads without a family and check the reliability of the call, which increases substantially the sequencing depth for variant calling, a particular important advantage for low-input samples or low-coverage regions.

## Introduction

The identification of ultra-rare variants has been relevant in a range of diverse fields such as cancer research, tumor development and residual disease, somatic mosaicism, evolutionary biology, and epidemiology (reviewed in (Salk, Schmitt, and Loeb 2018)). As such, the last decade has seen an extensive development of technologies for the identification of variants occurring at very low levels (10^−4^ to 10^−9^). Different next generation sequencing (NGS) protocols based on short paired-end reads (150-300 nucleotides) have been developed for this purpose. To overcome the high error-rates (0.1-2%) associated with this sequencing platform (Schmitt et al. 2015), different approaches for library preparation have been published that include the addition of tags during library preparation either by a random sequence in the amplification primers (Jabara et al. 2011; Schmitt et al. 2012) or the hybridization of indexed molecular inversion probes (MIPs) (O’Roak et al. 2012; Hiatt et al. 2013). The common strategy of these approaches is that reads are grouped into families, out of which a consensus sequence is built. Real substitutions present in the majority of the reads of a family can be distinguished from PCR and sequencing errors (Jabara et al. 2011; O’Roak et al. 2012; Schmitt et al. 2012; Hiatt et al. 2013). Alternatively, family members can be created by the circularization of small DNA fragments followed by rolling circle amplification (Lou et al. 2013).

This grouping strategy reduces error rates to less than 10^−5^; however, amplifiable DNA lesions (such as 8-oxoguanine, or deaminated cytosine or 5-methylcytosine) affect the detection limits because they cannot be distinguished from true variants (Arbeithuber et al. 2016). This is resolved in duplex sequencing (Schmitt et al. 2012; Kennedy et al. 2013; Ahn et al. 2015), a strategy that tags both strands of the DNA by the ligation of adapters with a random barcode (Fig 1). The paired-end reads are then grouped into families or single strand-consensus sequences (SSCS) representing the forward (*ab*-SSCS) or reverse (*ba*-SSCS) strand that are then re-united into the original duplex consensus sequence (DCS). While a substitution is present in both DNA strands, errors due to DNA lesions are only found in one DNA strand and can be distinguished in duplex sequencing (DS) (Arbeithuber et al. 2016), making DS currently the method with the lowest error rate (Schmitt et al. 2012; Schmitt et al. 2015; Salk et al. 2018).

**Figure 1.**
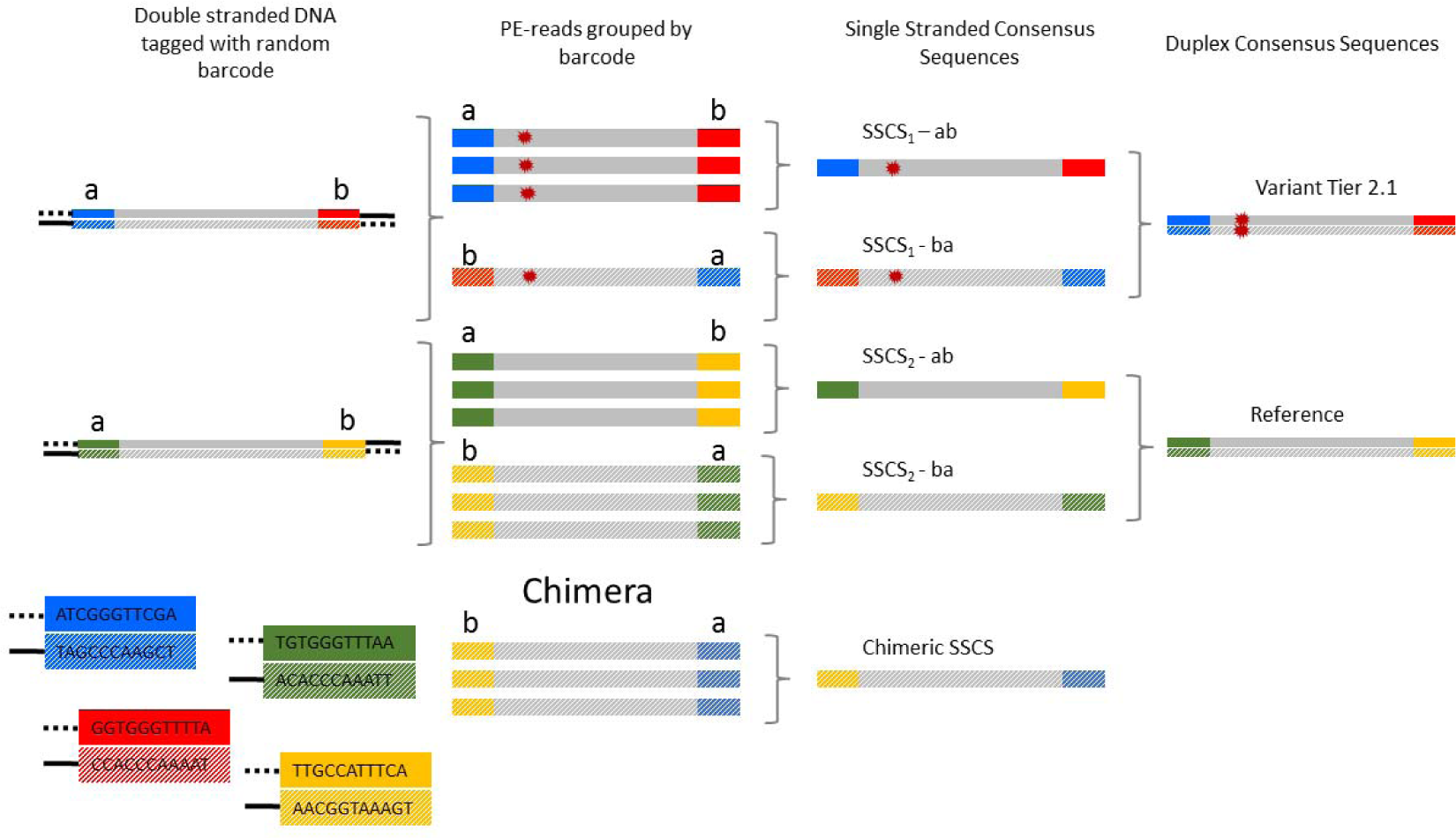
Schematic representation of duplex sequencing. An adaptor with a unique barcode is ligated to each end (*a* or *b*) of a double stranded DNA molecule (adaptors contain 10 random base pairs as exemplified in the red, green, blue, and yellow adaptors). During library preparation multiple copies are created from the original DNA by PCR that are then sequenced. PE-reads with the same barcode in either *ab* or *ba* orientation are grouped together into a family or single stranded consensus sequence (SSCS). Complementary SSCS (e.g. SSCS_1_-*ab* + SSCS_1_-*ba*) are united into a duplex consensus sequence (DCS) and substitutions found in both complementary SSCS are classified as true variants. Usually, a minimum of three reads per SSCS are required for consensus building and SSCS with single reads are discarded (e.g. SSCS_1_-*ba*). In our analysis, we include these and validate the variant calling by a tier-based system. Chimeras form by template switching or by incomplete extensions in PCR and are identified by sharing an identical *a* or *b* end with another SSCS family. These chimeric SSCS will unlikely have a complementary partner and might not form a DCS.

However, duplex sequencing is still quite costly and only a fraction of the input material (1%) results in a duplex consensus sequence (Schmitt et al. 2015; Nachmanson et al. 2018). Changes to the library preparation, such as CRISPR/Cas targeted digestion, reduces the number of amplification steps and increases the number of DCS per input material (6-12%) (Nachmanson et al. 2018). Also changes in the bioinformatic pipeline have improved the data output such as the Du Novo reference-free consensus building (Stoler et al. 2016) and error correction in the tags to re-unite otherwise “lost” reads into their respective families (Stoler et al. 2018).

Yet, there is still room for improvement, but this requires a better understanding of the sequencing data produced in DS. Moreover, tools for building DCS require a series of decision taking steps to ensure a low number of false positives. Quality control tools can enable informed decisions about the parameter settings during consensus building. These include deciding on a minimal number of reads per family or SSCS, the proportion of reads within the family with the alternative base, the number of errors allowed in the tag, and length of sequence trimming and removal of low quality bases. Currently, most of these steps are performed automatically using a “one-fits-all” approach with settings that ensure that a low number of false positives or low quality consensus reads end up in the DCS. The downside is that with a very conservative setting, more data is lost.

Here we created a series of quality control (QC) tools (Tag Distance, Chimera Analysis, Family Size Distribution, and Variant Analyzer) that can be implemented at different steps of consensus building and have been tested within Du Novo (Stoler et al. 2016) and compared against the output obtained with the software developed by (Nachmanson et al. 2018). The purpose of these tools is: 1) Allow for an informed decision as to the best analysis parameters for a particular dataset (currently done by trial and error). 2) Minimize the number of false positives and false negatives, and at the same time, maximize the number of consensus calling (DCS). 3) Allow variant calling with more relaxed parameters in the consensus building steps since calls are re-evaluated by a series of summary data validating a variant by a tier-based system that can then be manually examined.

## Results and Discussion

### Datasets

In order to test our tools, we used a previously published dataset produced by (Nachmanson et al. 2018), who sequenced the *TP53* exonic regions with DS using the standard protocol (Schmitt et al. 2015) and a targeted genome fragmentation approach based on CRISPR/Cas9 digestion. In particular, we focused on the following four libraries: PF1-CRISPR, PF1-Standard, PF2-CRISPR, and PF2-Standard. The main difference between PF1-Standard and PF2-Standard is the starting amount of genomic DNA with ∼10 or 3 µg and the allele frequency of variant chr17-g-7674230C>T (∼68% vs ∼1% in PF1 and PF2, respectively). Note that the amount of starting material used in the PF1-CRISPR and PF2-CRISPR libraries was 100 ng of DNA.

### Quality control tools

Our QC-tools comprehend the following analysis: 1) Tag Distance (TD), 2) Tag Distance (TD) with the Chimera Analysis (CA), 3) Family Size Distribution (FSD), and 4) Variant Analyzer (VAR-A). The TD, CA, and FSD tools analyze the tag composition extracted from the paired-end reads (PE-reads) and are useful for deciding on parameter settings in the bioinformatics pipeline. The tag composition also provides important insights on how different library preparation protocols affect the yields of duplex consensus data. VAR-A provides a comprehensive summary of the called variants by re-analyzing the PE reads, such that the calling of rare variants is verified and borderline cases can be manually inspected. Here we present the results of the tools implemented in the user-friendly Galaxy environment following a general pipeline (Fig S1) that can be implemented as part of the DS analysis (see Supplemental Note 1).

### Tag distance (TD)

In DS, each paired-end sequencing read (PE-read) is tagged by an upstream (*a*) and downstream (*b*) random barcode resulting in a tag that labels either the forward (*ab*) or the reverse strand (*ba*) of the input DNA (see Fig 1). All the reads containing the same tag (either in the *ab* or *ba* orientation) are grouped into a family or single stranded consensus sequence (SSCS) for the forward (*ab*) or the reverse strand (*ba*). The duplex consensus sequence (DCS) is then formed by uniting complementary *ab*+*ba* SSCS. In the library preparation, a large excess of unique tag combinations (4^10+10^ or 1.1 × 10^12^) was used to label the DNA templates (e.g. 1-3 million templates). This large tag-to-template excess makes it very unlikely that highly similar barcodes are used. However, sequencing or PCR errors could introduce 1-2 nucleotide differences in the tags that would be otherwise identical. These tags with errors will be effectively lost if not re-united with a barcode correction tool to the original family (Stoler et al. 2018).

The TD tool analyzes this phenomenon by calculating the number of nucleotide differences among tags, also known as Hamming distance (Hamming 1950). The tool compares the number of sequence differences of a subset of 1000 tags with the rest of the tags in the dataset and plots the smallest number of differences with another tag (minimum tag distance) as a histogram stratified by family size (Fig 2). Note that a sample subset of 1000 tags comprehend ∼0.1% of the sample size; however, this sample size is representative given that more data (e.g. 1% or 10%) rendered very similar results, but was computationally more time-intensive (see Fig S2).

**Figure 2.**
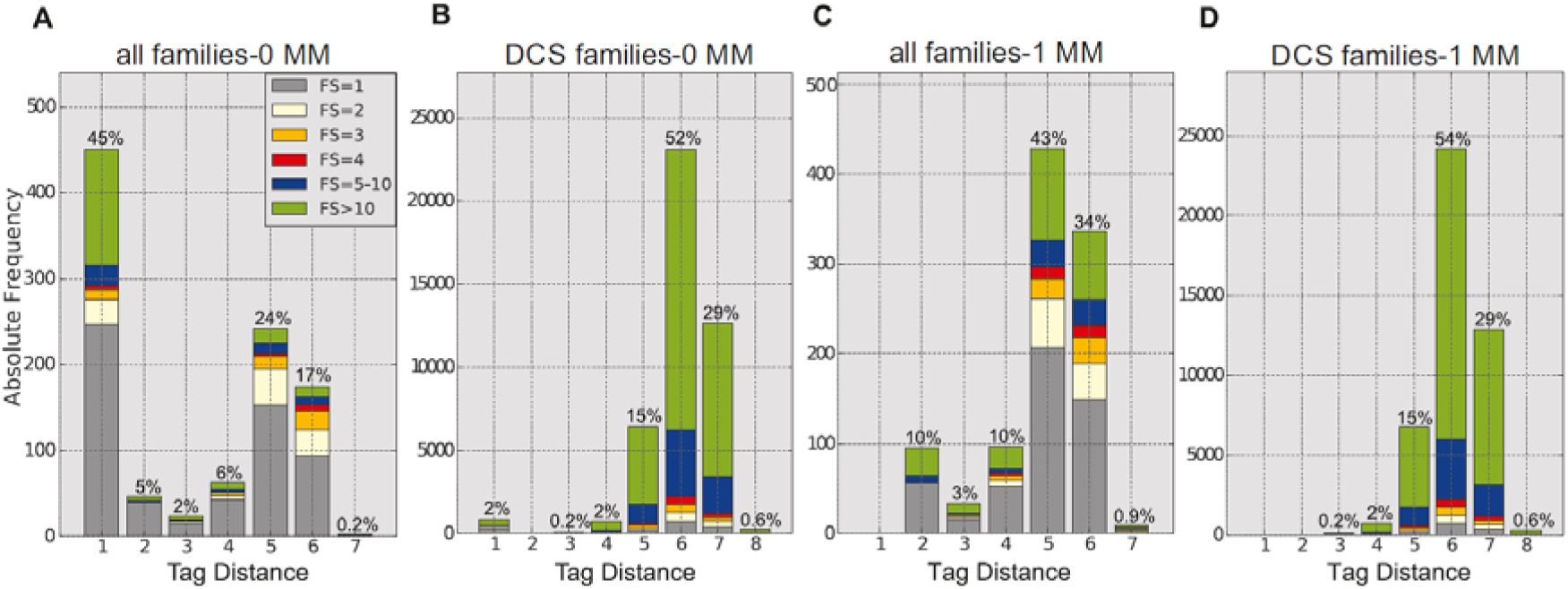
A) Tag distance (TD) before barcode correction in the PF2-CRISPR library. The TD was estimated for a subset of 1000 tags, each tag representing a family or SSCS. The legend denotes the family size (FS)-number of reads with the same tag grouped into a family **B)** TD distribution using tags that can form a DCS (complementary *ab*+*ba* tags). Note that in this case all tags in the DCS dataset were used for the estimation of the TD. **C**) TD distribution after barcode correction allowing one mismatch in the tag. TDs smaller or equal to the implemented barcode correction parameter are now reduced to zero. **D)** TD distribution after barcode correction using tags that form DCS.

In the PF2-CRISPR and PF2-Standard data sets, the length of the random barcode is 10 nucleotides on each side of the PE-read (10+10). The TD analysis of these data sets shows a bi-modal distribution of the tag distance (Fig 2A and Fig S3A): a little less than half of the tags differ by 5-7 nucleotides (TD=5-7), while 30-45% of the tags differ only by one nucleotide (TD=1). In general, for bi-modal tag distributions like these, the smaller TD is indicative of sequencing or PCR errors in the tags. This is verified in a TD analysis using only tags forming DCS, which should be error-free (Fig 2B and Fig S3B show that almost no tags with TD=1 are found in the DCS population). Thus, in these data sets it is appropriate to implement the barcode correction tool (Stoler et al. 2018) allowing for at least one mismatch in the tags, which improves the duplex coverage from 23% to 32% and 18% to 23% in the PF2-CRISPR and PF2-Standard libraries, respectively. As expected, the same analysis of TD after the barcode correction with one mismatch now renders mainly tags with a TD = 5-7, since tags with one difference have been re-united with larger families (thus, TD =1 is zero).

The TD tool is particularly useful to decide how many errors or mismatches (1, 2 or 3) can be allowed in a tag such that it is re-grouped with its family. This is an important aspect, especially for libraries prepared with less barcode combinations (e.g. shorter random barcodes of 8 or 6 nucleotides). The shorter the tag, the higher the frequency of tags with one or two differences that cannot be distinguished from sequencing or PCR mistakes. This is illustrated in Fig 3, in which we computationally shortened the tag from 10+10, to 8+8, and 6+6 nucleotides. As tags get shorter, the proportion of tags with a TD = 4-7 decreases and the proportion of tags with small distances increases (see Table 1). In short tags (e.g. 6+6), errors cannot not be distinguished anymore from real differences between uniquely labelled molecules.

**Table 1.** Proportion of tags with the given tag distance (TD) when computationally shortening the length of the original tag (10+10 nt) obtained in the PF2-CRISPR library. The number of total tags represents all the different families/tags in the dataset.

| tag length (nt) | TD = 1 | TD = 2-3 | TD = 4-7 | Total tags |
| --- | --- | --- | --- | --- |
| 20 (10+10) | 45% | 6.9% | 48.1% | 188354 |
| 16 (8+8) | 43.3% | 31.6% | 25.1% |  |
| 12 (6+6) | 57.7% | 42.3% | 0% |  |

**Figure 3.**
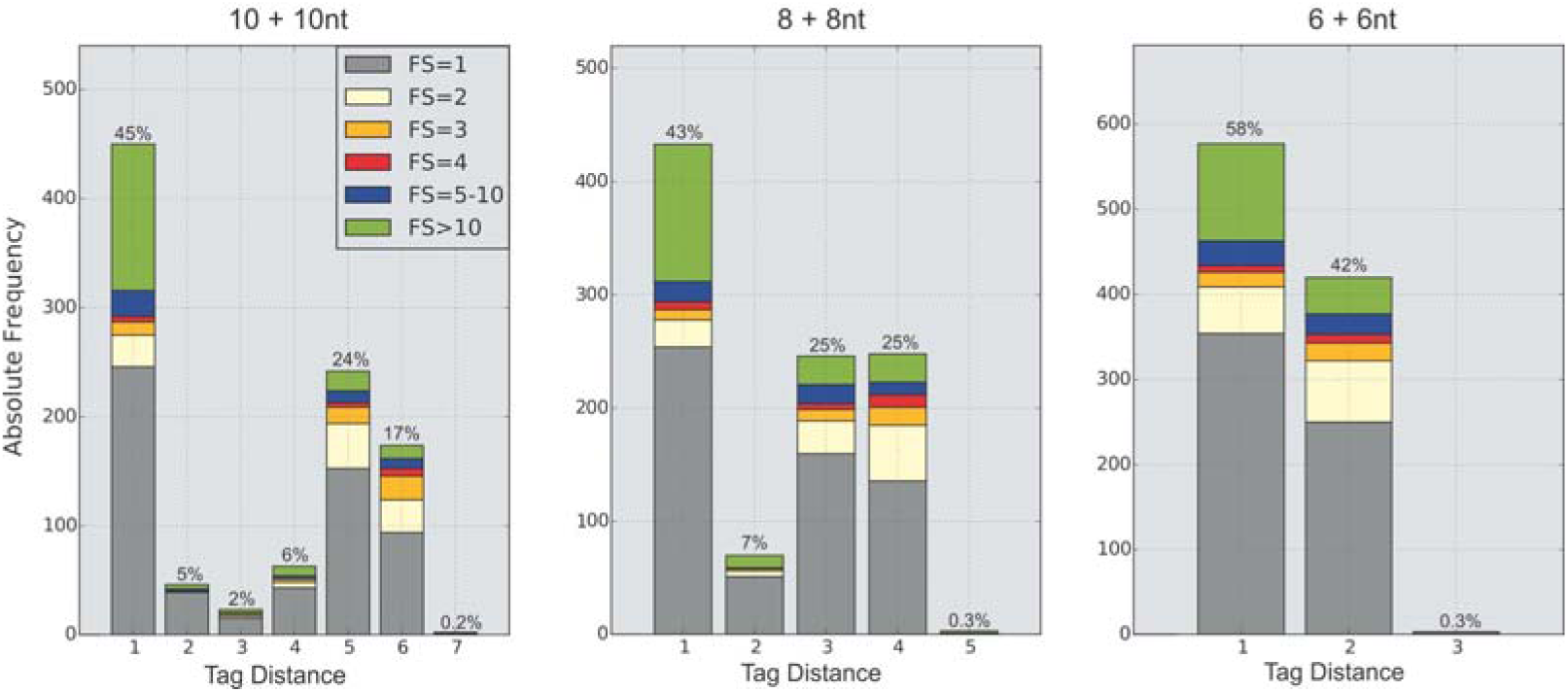
Effect of computationally shortening the length of the original tag in PF2-CRISPR. The tag (10+10) is shortened to 8+8, and 6+6 nucleotides after splitting the full tag (20 nucleotides) in half and removing 1 and 2 nucleotides at both ends of the half.

### Chimera Analysis (CA)

Chimeric reads are a known problem in PCR-based methods when a primer is not completely extended during PCR and this partial extension acts as a primer on similar templates, or by template switching in which an extended strand jumps from the original template to another one (Odelberg et al. 1995; Kanagawa 2003). During PCR, chimeras form at a frequency of 0.2% or 20% (if the plateau phase is reached in PCR), regardless of the proof-reading activity or processivity of the PCR polymerase (Boulanger et al. 2012). The formation of chimeras could be a serious problem in DS, because it confounds copies derived from the same template as two independent copies, and potentially leads to a false estimate of variant frequencies. In addition, chimeras reduce the DCS coverage since chimeric SSCS will unlikley have a complement to form a duplex.

We have developed the *Chimera Analysis (CA) based* on the examination of differences between the tags at both ends of a read (*a/b*). Chimeras can be distinguished by carrying the same tag at one end combined with multiple different tags at the other end of a read (Fig 1). Given the large excess of random barcodes to input templates, it is very unlikely that two identical tags are being used to label different templates more than once. The CA works as an extension of the TD tool and computes differences among tags in the data set. Basically, it searches tags that are identical at one end and different at the other end of the read. This is done by splitting the tag into two parts (*a* and *b*) in a subset of 1000 tags (each part representing the upstream/downstream end). Tags with the smallest distance in the *a* part (TD*a*_*min*_) and the largest distance in *b* (TD*b*_*max*_) in the data set (and vice-versa: smallest distance in *b* and largest distance in *a*) are extracted. It then analyzes the contribution of each part (*a* + *b*) to the overall TD. In a chimera, it is expected that only one end of the tag contributes to the TD of the whole tag. In other words, if the same *a* part is observed in combination with several different *b* parts (as would be expected in a chimera), then one end will have a TD=0. Thus, the TD difference between the parts (TD*a*_*min*_-TD*b*_*max*_) is the same as the sum of the parts (TD*a*_*min*_+ TD*b*_*max*_) or the ratio of the difference to the sum (*relative delta TD =* TD*a*_*min*_-TD*b*_*max*_ */* TD*a*_*min*_+ TD*b*_*max*_) will equal to one in chimeric families.

In the PF2-CRISPR or PF2-Standard libraries, ∼44% or 47% of the tags are formed by chimeric families with a *relative delta TD* value of one (Fig 4A and Fig S4A), respectively. Interestingly, the majority (80%) of these chimeras just form SSCS, but not DCS (Fig 4B and Fig S4B). An important source of data loss in DS, are SSCS than cannot be grouped into DCS. It is possible that these unpaired SSCS are the result of an amplification bias, in which only the forward or the reverse strand gets amplified, as is the case in ∼ 20-50% of the templates in PCR (see this phenomenon in (Arbeithuber et al. 2016)). Another source of unpaired SSCS could be chimeras, since it is highly unlikely that chimeric SSCS will have a complement and form DCS. In fact, 8% of the DCS are formed by chimeric families (Fig 4C and Fig S4C). Note that the DCS chimeras are mainly coming from very large families with more than 10 members (Fig 4D and Fig S4D), suggesting that this phenomenon is a PCR artefact. Eliminating chimeras could be an important modification in the library preparation protocols to improve DCS coverage. This could be achieved by avoiding amplifying mixtures of very similar DNA, but instead using a single molecule format using for example a water/oil emulsion as described previously (Palzenberger et al. 2017).

**Figure 4.**
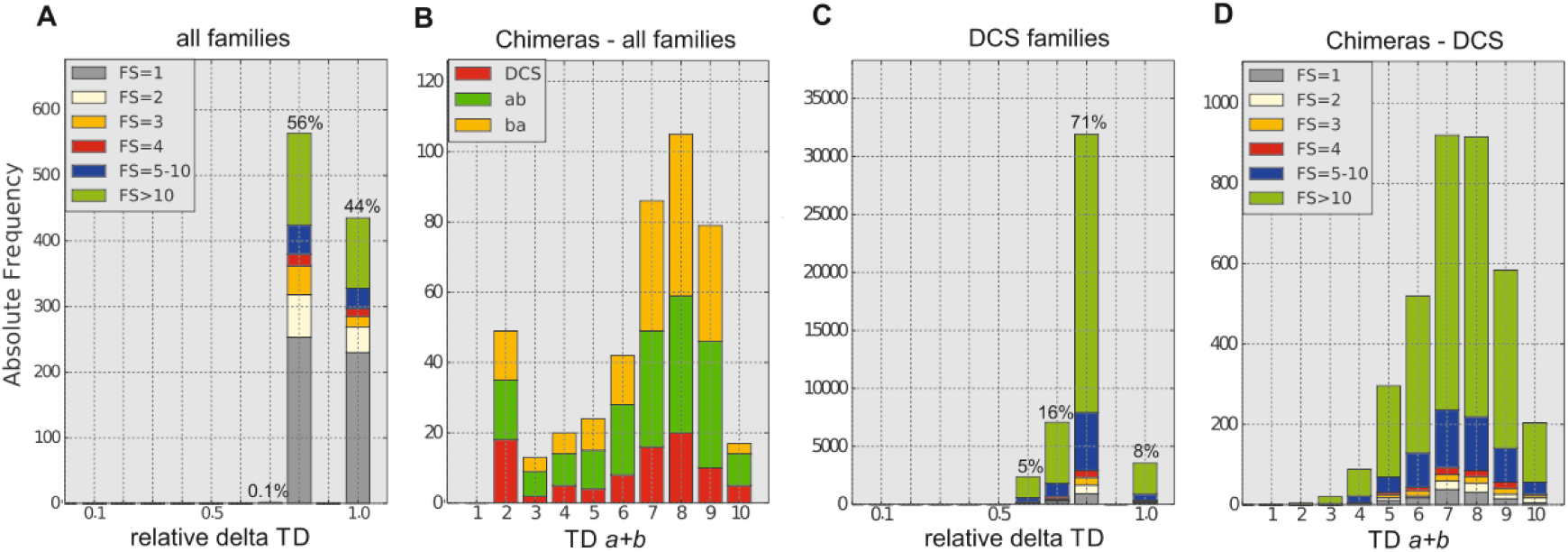
**A.** Chimera analysis (CA) of a subset of tags (n=1000) derived from the SSCS of the PF2-CRISPR library after barcode correction (allowing one mismatch). Chimeras have a *relative delta TD*=1. **B**. Distribution of TD in chimeric tags (with a *relative delta TD*=1) stratified into DCS or SSCS. Out of 435 chimeric reads, 181 (42%) belong to SSCS ab; 166 (38%) to SSCS ba, and 88 (20%) to DCS. **C**. Chimera analysis using the tags of DCS and **D**. tag distance of the chimeric DCS. Note that most of the chimeric DCS are from large family sizes (>10).

### Family size distribution (FSD)

One of the first steps in the consensus building in duplex sequencing is grouping together PE-reads representing copies of an initial molecule into a family or consensus (in this case SSCS). The more reads within a family carrying the substitution, the more likely the substitution is real and not an artefact. There is a delicate balance around the optimal family size: small families make variant calling less reliable, while larger families reduce the DCS coverage and total yields. Our FSD tool analyzes the family size associated with each tag and renders a graphical and tabular output of the absolute and relative family sizes compared to the total amount of families and total amount of PE-reads. This tool can be used to compare the FSD among different libraries or different steps of the bioinformatic pipeline (e.g. barcode correction or sequence trimming).

This tool also analyzes the ratio of SSCS/DCS for each family size. This latter analysis (Fig 5) is particularly useful to decide on the minimal number of PE-reads to build a consensus sequence (SSCS). In the early days of DS, an average family size of 6-7 members was considered appropriate for reliable variant calling (M. W. Schmitt et al. 2012), but with time this number has been reduced to three members to increase the data yields and DCS coverage (Nachmanson et al. 2018). The decision as to the minimal number of PE-reads to form a consensus depends on each library, thus having a tool to visualize the family size distribution is quite useful. In the PF2-CRISPR and PF2-Standard libraries, 32% or 23% of the families are united into DCS (68% or 33% of the total PE reads), respectively, and more than half of the DCS are formed by family sizes between 3 and 20 members (Table 2, Fig 5). Thus, for these libraries using a family size with a minimal number of three reads is recommended (Nachmanson et al. 2018); although, with this setting 5-12% of the DCS, formed by smaller family sizes of 1-2 reads, get lost.

**Table 2.** Fraction of data forming duplex consensus sequences (DCS) from the total number of families or PE-reads obtained in each library (column 1 and 2, respectively). We counted 31710, 147944, 44912, and 253428 families that can form a DCS for PF1-CRISPR, PF1-Standard, PF2-CRISPR, and PF2-Standard, respectively. Column 3-5 represent the proportion of DCS with a family size (FS) of 1-2 reads, 3-20 reads or more than 20 reads, respectively. Column 6 reports the average family size (FS) and column 7 shows the total number of families counted including single stranded consensus sequences (SSCS) and duplex consensus sequences (DCS).

|  | DCS families | DCS PE-reads | DCS FS1-2 | DCS FS3-20 | DCS FS>20 | mean FS | total families (DCS +SSCS) | total PE-reads |
| --- | --- | --- | --- | --- | --- | --- | --- | --- |
| PF1-CRISPR | 17% | 53% | 11% | 62% | 27% | 5 | 189,675 | 1,004,941 |
| PF1-Standard | 11% | 24% | 5% | 12% | 83% | 27 | 1,341,763 | 35,748,407 |
| PF2-CRISPR | 32% | 68% | 5% | 56% | 39% | 9 | 138,631 | 1,284,504 |
| PF2-Standard | 23% | 33% | 12% | 64% | 23% | 11 | 1,106,303 | 11,717,544 |

**Figure 5.**
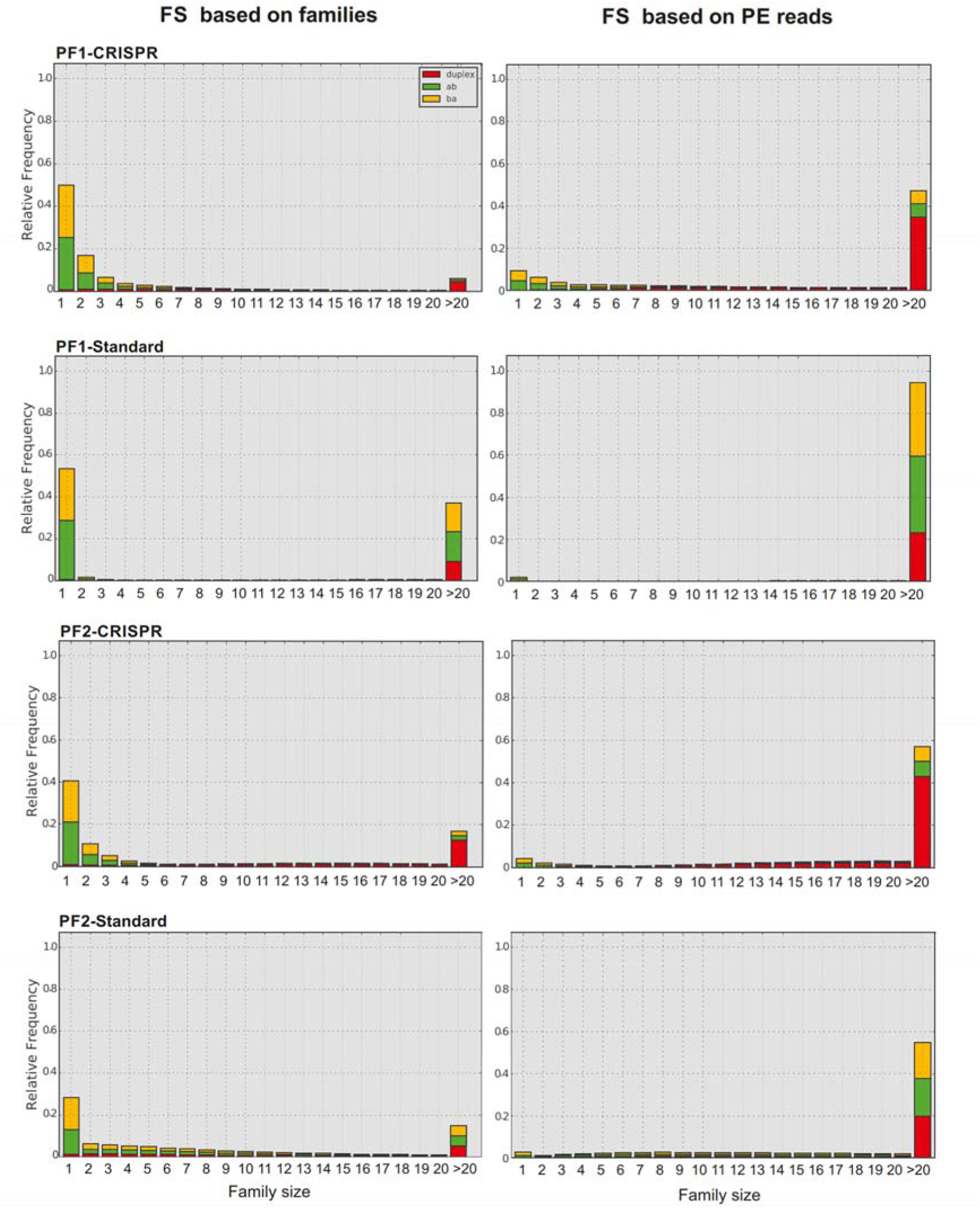
Family Size (FS) distribution in the different libraries. Frequency of reads per family observed in the four analyzed libraries (PF1-CRISPR, PF-1 Standard, PF2-CRISPR, and PF2-Standard) relative to the total number of families or to the total number of PE reads. Family size (FS) is stratified into SSCS (*ab* or *ba*) or DCS (*duplex*).

Knowing the ratio of SSCS/DCS for each family size also helps to understand the data allocation and DCS yields: for example, Nachmanson and colleagues reported that CRISPR-DS is superior over standard-DS, given the 10-fold higher recovery rate of specific target regions (average DCS coverage per input templates) of the former (Nachmanson et al. 2018). Our FSD-tool provides a different light on the performance of these two DS protocols: in terms of DCS recovery and allocation of DCS within optimal family sizes (3-20 reads), only one of the four libraries (PF1-Standard) performed suboptimal (Figure 54, Table 2). In this library, ∼83% of the families (or ∼94% of the total PE-reads) had more than 20 reads, which explains the lower DCS coverage of this library. However, the other standard DS library (PF2-Standard), which used a third less of DNA, performed equally well than the two CRISPR libraries: all three libraries had 56-62% DCS formed by optimally sized families (3-20). With our tool, it can also be observed that PF1-CRISPR and PF2-Standard formed DCS with small family sizes (FS = 1-2), which would get lost if at least three reads are required in a family. Understanding what factors influence the formation of sub-optimal family sizes for DCS (very small or very large families) during library preparation is an intensive research focus and our FSD-tool supports a detailed evaluation.

### Variant Analyzer (VAR-A)

Currently in DS, a family or consensus (SSCS) is built with a minimum of three or more reads (FS≥3). Yet, 5-12% of the DCS in the four tested libraries were formed by families with 1-2 reads. These DCS are discarded; although, they might contain important variant information and their inclusion would increase the coverage. The reason for requiring larger family sizes is that small families bear a higher risk of false positive calls. We developed the Variant Analyzer (VAR-A) to assess the evidence supporting a variant call based on a series of different summary data extracted from the raw PE-reads that classifies the confidence level of a variant call by a tier-based system. This allows using more relaxed analysis parameters during consensus building, e.g. small families (including families with only one or two reads) or *ad hoc* stringent trimming parameters.

Only DCS with a variant in the original consensus building output are re-analyzed in VAR-A. The evidence of a variant is then compiled from the raw PE-reads, and includes information on the mate (if both mates overlap the position of the variant), the number of high quality PE-reads in both forward (*ab*) or reverse (*ba*) SSCS, the proportion of alternate vs. reference calls within the family, and the median position of the variant within the consensus sequence. The quality of the variant call is re-analyzed, PE-reads are trimmed automatically, and positions with a low PHRED score are removed. Given these additional analysis layers, more data can be used for consensus calling without compromising the reliability of the analysis and false negatives can get potentially recovered. The tabular output of VAR-A also includes relevant information such as the sequence of the tag, if more than one variant is present in a family (multiple consecutive variants in one molecule), and if the variant is part of a chimeric family.

VAR-A also categorizes variants with a tier-based system (see Table 5 and Table S1) that helps the user to distinguish high quality calls from those with lower support. Tier 1 variants have the strongest support with information from multiple reads and mates. However, the inclusion of second order tiers (1.2-2.4) increases the coverage without seriously compromising the accuracy or reliability of the call that can be removed if necessary after manual inspection. These second-order tiers are particularly interesting, because they include small families (1-2 reads) for either the forward or the reverse SSCS, which would be discarded by the regular pipeline. Yet, these variants are likely real since they are present in both forward and reverse SSCS, albeit in one of them at a low number. Table 3 compares the variants identified using different settings for the minimum family size (FS≥3 or FS≥1). When reducing the minimum family size from three to one read, we rescued ∼2200 or ∼25,500 DCS resulting in a ∼10% or 25% increase of DCS coverage in the PF2-CRISPR or PF-2 Standard library, respectively (see Table 3).

**Table 3:** shows the coverage of the Du Novo analysis from the PF2-CRISPR and PF2-Standard library using different parameters for consensus building for the minimum family size (FS). Reducing the FS settings to FS≥1 increased the number of DCS.

| **PF2-CRISPR** | **FS $\geq 3$** | **FS $\geq 1$** | **PF2-Standard** | **FS $\geq 3$** | **FS $\geq 1$** |
| --- | --- | --- | --- | --- | --- |
| **Total reads** | 1,284,504 |  | **Total reads** | 11,717,544 |  |
| **Total SSCS** | 138,631 |  | **Total SSCS** | 1,106,303 |  |
| **SSCS FS 1-2** | 116,175 |  | **SSCS FS 1-2** | 979,589 |  |
| **DCS** | 20,218 | 22,456 | **DCS** | 101,145 | 126,714 |
| **On target** | 20,217 | 22,426 | **On target** | 101,095 | 125,826 |

**Table 4.**
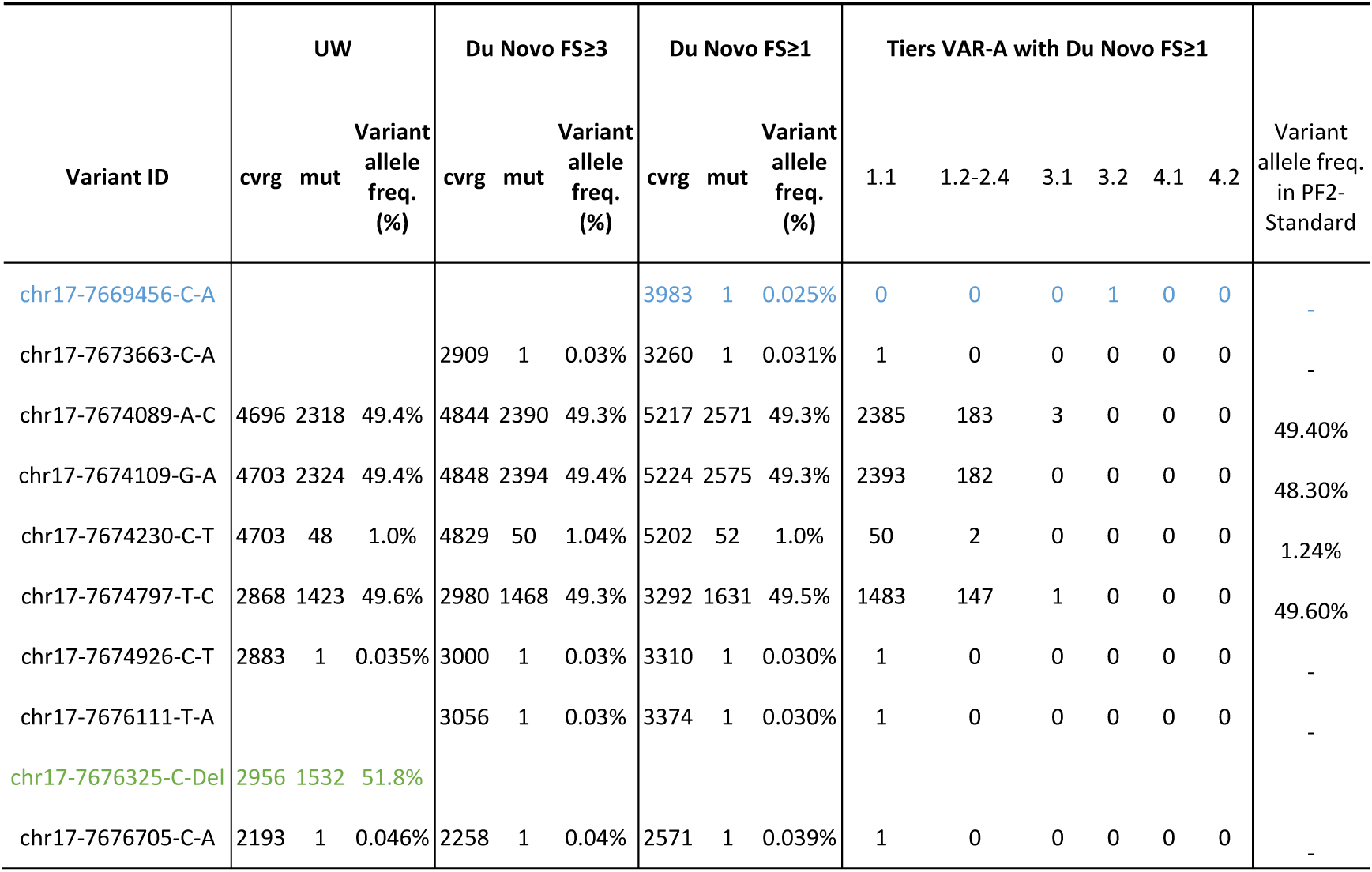
Summary of the variants identified in the PF2-CRISPR data set. We compared the original pipeline of University of Washington (UW) with Du Novo analysis using FS ≥ 3 or FS ≥ 1, the latter combined with VAR-A. The variants identified in VAR-A are classified by the different tiers representing high or lower confidence calls. Details of variants are shown in Table S3. The variant ID is based on the human genome assembly GRCh38/hg38. Variants in blue are only detected by the Du Novo run with relaxed settings such as a family size (FS ≥ 1). Variants in green were detected only by the pipeline of UW. Note that this is due to the fact that VAR-A currently does not support the evaluation of in-dels.

**Table 5.** Definition of the tier system in the Variant Analyzer tool. Tiers 1.2-2.4 include SSCS with small family sizes (<3) and would be discarded by the regular pipeline, yet variants are likely real and are confirmed by both forward (*ab*) and reverse (*ba*) SSCS.

|  |  |
| --- | --- |
| Tier 1.1 | both *ab* and *ba* SSCS present (>75% of the sites with alternative base) and minimal FS≥3 for both SSCS in at least one mate |
| Tier 1.2 | both *ab* and *ba* SSCS present (>75% of the sites with alt. base) and mate pair validation (min. FS=1) and minimal FS≥3 for at least one of the SSCS |
| Tier 2.1 | both *ab* and *ba* SSCS present (>75% of the sites with alt. base) and minimal FS≥3 for at least one of the SSCS in at least one mate |
| Tier 2.2 | both *ab* and *ba* SSCS present (>75% of the sites with alt. base) and mate pair validation (min. FS=1) |
| Tier 2.3 | both *ab* and *ba* SSCS present (>75% of the sites with alt. base) and minimal FS=1 for both SSCS in one mate and minimal FS≥3 for at least one of the SSCS in the other mate |
| Tier 2.4 | both *ab* and *ba* SSCS present (>75% of the sites with alt. base) and minimal FS=1 for both SSCS in at least one mate |
| Tier 3.1 | both *ab* and *ba* SSCS present (>50% of the sites with alt. base) and recurring mutation on this position |
| Tier 3.2 | both *ab* and *ba* SSCS present (>50% of the sites with alt. base) and minimal FS≥1 for both SSCS in at least one mate |
| Tier 4.1 | variants at the start or end of the DCS (median position of the variant within the PE-reads forming the DCS) |
| Tier 4.2 | other |

Since small families make variant calling less reliable, we analyzed the variants with VAR-A using large (FS ≥ 3) or small family sizes (FS ≥ 1) and compared them with variants identified with the bioinformatics pipeline from University of Washington (Nachmanson et al. 2018). The full output of the VAR-A analysis for the PF2-CRISPR and PF2-Standard data set is included as Table S3, Table S4, respectively.

As expected, heterozygous positions (50%) shown in Table 4 (PF2-CRISPR) or Table S2 (PF2-Standard) are detected by all three analyses at very similar frequencies while the coverage in the VAR-A analysis with FS ≥ 1 is higher, demonstrating that reducing FS ≥ 1 renders the same reliable calls as the other two more conservative pipelines, but at a higher coverage. While the increase in coverage might not seem extremely important in this example, it is especially an advantage in low-input samples, where an increase in sequencing depth for variant calling is crucial.

With VAR-A, we also identified important biases or false positives in the DCS of a library. For example, note that in the standard library (Table S2 and Table S4) some variants occur with a low tier (mainly 4.1-4.2). Some of these variants are of high sequence quality and form large family sizes, but co-occur at the end or beginning of the DCS, either as single independent events or as multiple variants within the same family (e.g. chr17-7676340-G-C and chr17-7676341-T-C). This strong positional bias of variants (mainly at the beginning of the DCS, representing the 5’ or 3’ end of the original DNA molecule) makes it highly likely that these are the result of end-polishing during A-tailing before adapter ligation. Variants at the beginning or the end of DCS are labelled by VAR-A as tier 4.1 and can be manually discarded. Another example is variant chr17-7675393-C-T that occurs close to a poly-T homopolymer (chr17-7675394-7675411) reported to be associated with noisy reads (Nachmanson et al. 2018) and that was tagged by VAR-A as a low tier that can be manually inspected and discarded.

When building consensus sequences with a FS ≥ 1, VAR-A validated two new variants that were missed by the analysis using a FS ≥ 3 (shown in blue in Table 4 and Table S2). These occur at an ultra-low frequency (∼10^−4^), which underlines the power of VAR-A in terms of detecting very rare variants. The identified variant (chr17-7669456C-A) in the PF2-CRISPR library classified as tier 3.1 is formed by SSCS with 3 and 2 members; in one SSCS, two out of three reads carried the alternative allele and in the other SSCS both reads carried the variant. This call could be considered a borderline case and would require further validation in subsequent libraries with higher coverage. We did not identify this variant in PF2-Standard (DS of the same biological sample) likely because of insufficient coverage in this library at this position. Similarly, in the PF2-Standard library, variant chr17-7675381T-C was counted twice (with tier 2.1 and 2.4) totaling to an AF of 0.02%. Further inspection of this variant showed that it is formed by high quality PE-reads including tier 2.1, 2.4 and tier 4.1 (the latter representing the end of a DCS) shown in Table S5. This variant also could be a borderline case since it occurs close to the poly-T homopolymer (chr17-7675394-7675411). Note that most of the variants in these positions were filtered out during the different QC steps of VAR-A (tier 4.2), as were those calls close to another poly-T homopolymer (chr17:7674361-7674376).

We also obtained the same variant allele frequency (1.0% or 1.2% for the PF2-CRISPR or PF2-Std, respectively) for variant chr17-7674230C-T with the relaxed pipeline as reported by Nachmanson and colleagues (labelled as chr17-7577548C-T with GRCh37/hg19 as a reference) (Nachmanson et al. 2018). More interestingly, performing the analysis with FS≥ 1 increased the coverage of this low frequency variant by ∼9% or ∼42% (from 4829 to 5202 DCS or 2003 to 3463 DCS in the PF2-CRISPR or PF2-Standard library, respectively). This example highlights the value of VAR-A to accurately call low variant frequencies at a quite improved coverage. Note that VAR-A also adds relevant biological information such as the haplotype associated with the mutation or if a mutation is part of a chimeric family.

We conclude that VAR-A is a powerful tool to increase the coverage without compromising the reliability of the variant calling, which is especially advantageous in low-input samples or low-coverage regions, where an increase in sequencing depth for variant calling is of particular importance.

## Conclusions

The analysis of the tag composition with our tools has been an important aspect to better understand the sources of data loss during the bioinformatics analysis. Moreover, with our tools we show how many reads are part of very large families or are singleton reads (without a family) and contribute to data loss and lower DCS coverage and yields. We also show that chimeras result in unpaired SSCS and reduce DCS yields and could introduce noise in the estimation of rare variant frequencies by inflating wild type or variant numbers. Finally, we demonstrate that reads with small families can be included in the consensus calling since the quality and reliability is validated with our Variant Analyzer tool. The resulting increase in coverage is especially an advantage in low-input samples, where an increase in sequencing depth for variant calling is crucial and allows for an identification of ultra-low frequency variants without blindly increasing the risk of false positive calls.

## Methods

### Family size distribution (FSD)

We first trimmed the barcodes from all sequencing reads generating a list of 10 + 10 barcode combinations that were arranged in lexicographic order and then counted the number of times each combination appeared in this list (family size). Each tag represents a family of paired-end sequences forming SSCS. The FSD analyzes then the family size associated with a tag, that is the number of reads per tag and produces several histograms with the distributed family sizes.

### Tag distance (TD) analysis

The tag analysis distance was estimated as reported in (Stoler et al. 2018). We used the same list of tags as already described in the FSD. Since the datasets contained more than one million tags, the comparison of all tags was computationally too demanding. Thus, we parallelized the algorithm and selected 1,000 random tags from the data set and compared them to the whole dataset (189,675, 1,341,763, 138,631 and 1,106,303 tags after barcode correction for the PF1-CRISPR, PF1-Standard, PF2-CRISPR, and PF2-Standard datasets, respectively). For the DCS tags, the smaller sizes made it possible to use the complete data set (15,855, 73,972, 22,456 and 126,714 tags after barcode correction, respectively). We have verified that a sample of 1,000 tags (0.1% of the data) is representative for the whole dataset (see Fig S1). At each comparison we calculated the number of differences (tag distance) and reported only the smallest number of differences (minimum tag distance) observed with any other tag. The distances between tags were calculated using equation 1, where D_i,j_ is the number of sites where X_i_ and X_j_ do not match, k is the index of the respective site out of the total number of sites *n*. A detailed guideline for using the TD analysis is described in **Supplemental Note 1**.

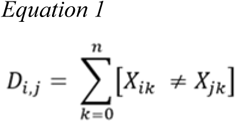

The output of the tool is a plot of the minimum tag distance (smallest number of differences) between tags as a frequency histogram categorized after the family sizes.

### Chimera analysis (CA)

We have extended the TD tool by the so-called *Chimera Analysis* which allows the identification of chimeric families in the sequencing data. Here, the analysis is described only for one tag in detail, but we repeat this process for 1,000 tags (default sample size for TD analysis). First, we split the tag into its upstream and downstream part (named *a* and *b*) and compare part *a* with all other *a* parts of the families in the dataset (∼1 million families). We estimate the sequence distance (TD) among the *a* parts and select those tags that have the smallest number of differences (TD*a*_*min*_) and then calculate from the subset the TD of the *b* part. The tags with the largest number of differences are extracted to estimate the maximum TD (TD*b*_*max*_). The process is repeated starting with the *b* part instead and estimates TD*a*_*max*_ and TD*b*_*min*_. Next, we calculate the absolute difference between TD*a*_*min*_ and TD*b*_*max*_ equal to *TD delta*.

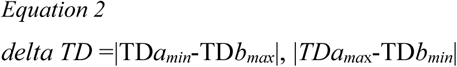

If the same *a* is observed in combination with several different *b* parts (as would be expected in a chimera), then *TD delta* will be large (low TD*a*_*min*_ and high TD*b*_*max*_). Thus, for chimeras, this difference is large, since the TD is contributed only by one of the parts (a or b), as the other part is identical (TD = 0). In order to normalize the values between comparisons, we use the relative *delta TD* defined as the ratio of *delta TD* to the sum of the TD of each part (TD a+ b).

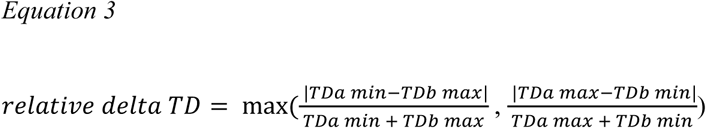

For chimeras, the larger of the two *relative delta TD* values is expected to be one, since only one part contributes to the TD. Note that error tag correction was performed before the chimera analysis. For more information on this analysis see **Supplemental Note 2**. The *Chimera Analysis* can also be used considering only tags that form DCS. A detailed guideline for using the CA is described in **Supplemental Note 1**.

### Variant Analyzer (VAR-A)

Du Novo’s analysis was performed with the following settings: barcode correction = 1 and family size = 1 and 3. This was followed by a trimming step (Blankenberg et al. 2010) of the 3’ and 5’ ends of the DCS with 10 nucleotides and the alignment to the human genome assembly GRCh38/hg38 using *BWA-MEM* (Li 2013) and *BamLeftAlignIndels* (Garrison and Marth 2012). Finally, we performed the variant calling with the *Naive Variant Caller (NVC)* (Blankenberg et al. 2014) and its annotation by the Galaxy tool *Variant Annotator*. Only variants with a minimum read depth of 100 reads were kept. A detailed workflow of the analysis can be seen in Fig S1. Each variant called by Du Novo with its subsequent analysis steps were re-analyzed by VAR-A. First, all tags of the DCS identified to carry a variant were extracted from the bam file obtained in the alignment step. Subsequently all PE-reads of these tags were extracted from the data and stored in a fastq file. Further, PE reads were trimmed using *Trimmomatic* (Bolger et al. 2014) with default settings and aligned to the reference using *BWA-MEM* (Li 2013). Finally, VAR-A outputs for each mutation, tag, mate, and direction (*ab*/*ba*) several statistics: e.g. the number and fraction of reads with reference and alternate allele, as well as the number of unaligned reads and reads (Phred-scaled base quality score < 20). If both mates overlap a variant, the second mate can either provide additional support for identifying a true variant or help to identify false positive calls. Therefore, confident variant calls become possible even if some of the families have very small sizes (1-2 reads). Furthermore, the output includes the median position of the variant within the reads and information about the number of SSCS carrying the variant, as well as, all other variants reported for the same tag. To help the user identify high from low confidence calls, we developed a tier system labelling each variant (Table 5, and Table S1).

## Data Access

The tools are written in Python, are open source and readily available through the user-friendly Galaxy platform that can be easily run without any programming experience https://usegalaxy.org/, and GitHub: https://github.com/monikaheinzl/duplexanalysis_galaxy and https://github.com/gpovysil/VariantAnalyzerGalaxy. All software is freely available under non-restrictive AFL 2.0 license.

The data was downloaded from https://www.ncbi.nlm.nih.gov/bioproject/?term=PRJNA412416 on January 23, 2019.

## Abbreviations

AF: allele frequency
SSCS: Single Strand Consensus Sequence
DCS: Duplex Consensus Sequence
DS: Duplex Sequencing
PCR: Polymerase Chain Reaction
FS: family size
RL: read length
TD: tag distance
PE: paired-end
CA: Chimera Analysis

## Disclosure Declaration

Ethics approval and consent to participate: Not applicable

Consent for publication: Not applicable

No Competing interests

## Funding

Funding for R.S., M.H., G.P., and I.T-B was provided by the Linz Institute of Technology (LIT213201001) and the Austrian Science Fund (FWFP30867000).

## Acknowledgements

We would like to thank Rosana Risques and Jesse Salk for providing the p53 dataset, Barbara Arbeithuber for advice on duplex sequencing and fruitful comments on the manuscript, and Ulrich Bodenhofer for help in the conceptualization stages of the tools.

## Authors’ contributions

GP and MH developed the software and performed the analyses, RS participated in the analysis and tested software components. NS improved and trouble-shooted software components. ITB conceived the project and ITB and AN provided funding, GP, MH and ITB wrote the manuscript.

## Notes

#### Summary of Updates

different abstract

https://github.com/monikaheinzl/duplexanalysis_galaxy

https://github.com/gpovysil/VariantAnalyzerGalaxy

https://www.ncbi.nlm.nih.gov/bioproject/?term=PRJNA412416

