## Supplementary Materials for "Increased yields of duplex sequencing data by a series of quality control tools"

**Running Title: Quality control in duplex sequencing data**

### Supplemental Materials

#### Supplemental Tables

**Supplemental Table S1.** Definition of the tier system in the Variant Analyzer tool. Tiers 1.2-2.4 include SSCS with family sizes ≥1 and would be discarded by the regular pipeline. Confirmation comes from both forward (*ab*) and reverse (*ba*) SSCS. Tier 4.1 are at the beginning or the end of a DCS and 4.2 are variants with very low quality scores that get discarded during the VAR-A analysis. We also included examples of VAR-A for the different tiers.

| Variant ID | DuNovo with FS≥3 | | | DuNovo with FS≥1 | | | Tiers VAR-A with DuNovo with FS≥1 | | | | | | Variant freq in PF2-CRISPR |
|---|---|---|---|---|---|---|---|---|---|---|---|---|---|
| | cvrg (tier 1.1-2.4) | mut (tier 1.1-2.4) | Variant fraction (%) | cvrg (tier 1.1-2.4) | mut (tier 1.1-2.4) | Variant fraction (%) | 1.1 | 1.2-2.4 | 3.1 | 3.2 | 4.1 | 4.2 | |
| chr17-7674089-A-C | 1133 | 544 | 48.0% | 2161 | 1067 | 49.4% | 576 | 491 | 0 | 0 | 14 | 0 | 49.30% |
| chr17-7674109-G-A | 1097 | 539 | 49.1% | 2017 | 974 | 48.3% | 547 | 427 | 0 | 0 | 23 | 0 | 49.30% |
| chr17-7674202-A-T | 1564 | 1 | 0.06% | 2842 | 1 | 0.04% | 1 | 0 | 0 | 0 | 0 | 0 | - |
| chr17-7674221-G-C | 1915 | 1 | 0.05% | 3368 | 1 | 0.03% | 1 | 0 | 0 | 0 | 0 | 0 | - |
| chr17-7674230-C-T | 2003 | 25 | 1.2% | 3463 | 43 | 1.24% | 25 | 18 | 0 | 0 | 0 | 0 | 1.00% |
| chr17-7674797-T-C | 2346 | 1154 | 49.2% | 2620 | 1299 | 49.6% | 1165 | 134 | 0 | 0 | 20 | 0 | 49.50% |
| chr17-7675019-C-T | 4187 | 1 | 0.02% | 4586 | 1 | 0.02% | 1 | 0 | 0 | 0 | 0 | 0 | - |
| chr17-7675327-C-T | 5612 | 2565 | 45.7% | 6355 | 3020 | 47.52% | 2549 | 471 | 0 | 0 | 28 | 0 | outside gRNA target |
| chr17-7675381-T-C | | | | 5286 | 2 | 0.04% | 0 | 2 | 7 | 0 | 12 | 33 | outside gRNA target |
| chr17-7675519-A-G | 1945 | 837 | 43.0% | 2601 | 1234 | 47.44% | 809 | 425 | 0 | 0 | 18 | 1 | outside gRNA target |
| chr17-7676435-G-T | 5333 | 1 | 0.02% | 6117 | 1 | 0.02% | 1 | 0 | 0 | 0 | 0 | 0 | - |

**Table S2. Summary of the variants identified in the PF2-Standard data set.** We compared the Du Novo analysis using FS ≥ 3 or FS ≥ 1 combined with VAR-A. Variants with only tier 3.1-4.2 were removed after manual inspection (shown in Table S4). The variant ID is based on the human genome assembly GRCh38/hg38. Variants in blue are only detected in the relaxed settings.

**Supplemental Table S2.** Summary of the variants identified in the PF2-Standard data set. We compared the Du Novo analysis using FS ≥ 3 or FS ≥ 1 combined with VAR-A. Variants with only tier 3.1-4.2 were removed (6 variants) after manual inspection (shown in Table S4). The variant ID is based on the human genome assembly GRCh38/hg38. Variants in blue are only detected in the relaxed settings.

**Supplemental Table S3.** Output of VAR-A for the PF2-CRISPR library relaxed analysis settings. The output contains the variant ID, which consists of the position of the variant compared to the reference (chr and pos) and the reference (ref) and alternate (alt) alleles. Next, the tier classification is listed as described in Table 5 and S1. The remaining columns contain the sequence of the tag, mate information, the median position within the reads (read pos.ab/read pos.ba), median length of the reads (read median length.ab/read median length.ba) and the median length of the DCS (DCS median length). Family sizes are reported before (FS.ab/FS.ba) and after QC (FSqc.ab/FSqc.ba), where QC means the removal of reads that could not be aligned to the reference (na.ab/na.ba) or had low PHRED scores at the position of the variant (lowq.ab/lowq/ba). In addition to the absolute number of reads with reference (ref.ab/ref.ba) or alternate (alt.ab/alt.ba), we also report their relative fraction based on the family sizes after QC (rel. ref.ab/rel. ref.ba/rel. alt.ab/rel. alt.ba). Next, we list the number of SSCS that carry the reference (SSCS ref.ab/SSCS ref.ba) or alternate alleles (SSCS alt.ab/SSCS alt.ba). If other variants were called within the same family, they are listed in column "other mut". The last column contains possible chimeric tags related to the tag analyzed.

**Supplemental Table S4.** Output of VAR-A for the PF2-Standard library relaxed analysis settings. The output format is the same as Table S3

#### Supplemental Figures

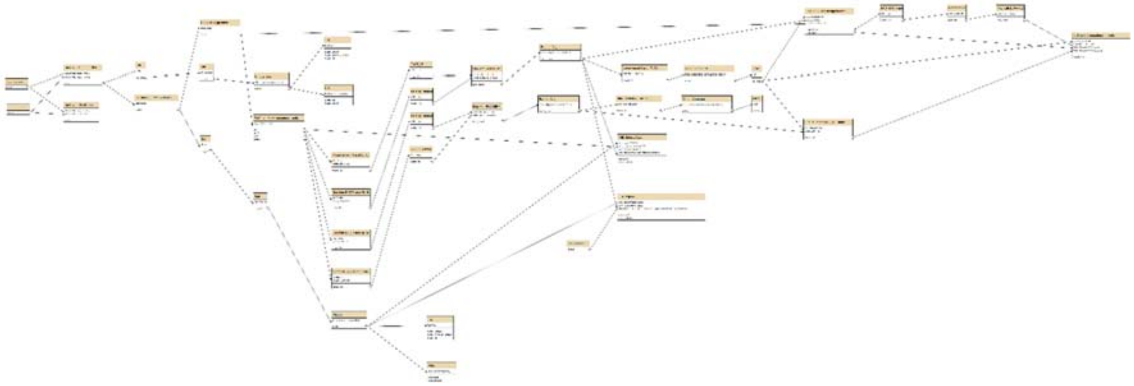

**Supplemental Figure S1.** Workflow of Du Novo and subsequent analysis steps in Galaxy system (larger version included separately) and can also be found under the link https://usegalaxy.org/u/jku-itb-lab/w/paperworkflow-du-novo-20.

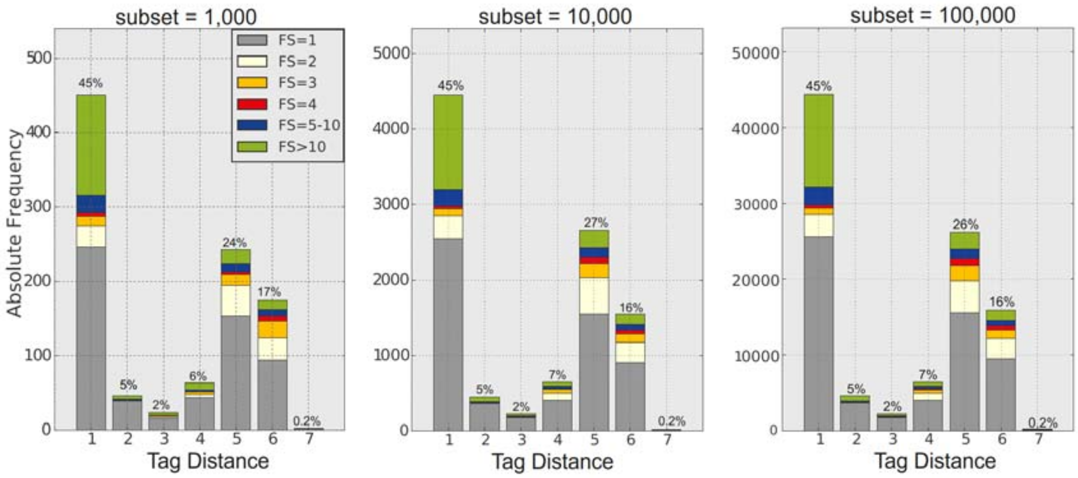

**Supplemental Figure S2.** Tag distance of PF2-CRISPR estimated in different sample subsets. Distribution of absolute numbers of tags with a given tag distance TD=1-7 using a subset of 1000; 10,000 or 100,000 tags. Note that barcode correction has not been implemented in these data. Data is stratified by family size (FS) and the data distribution is almost identical among datasets.

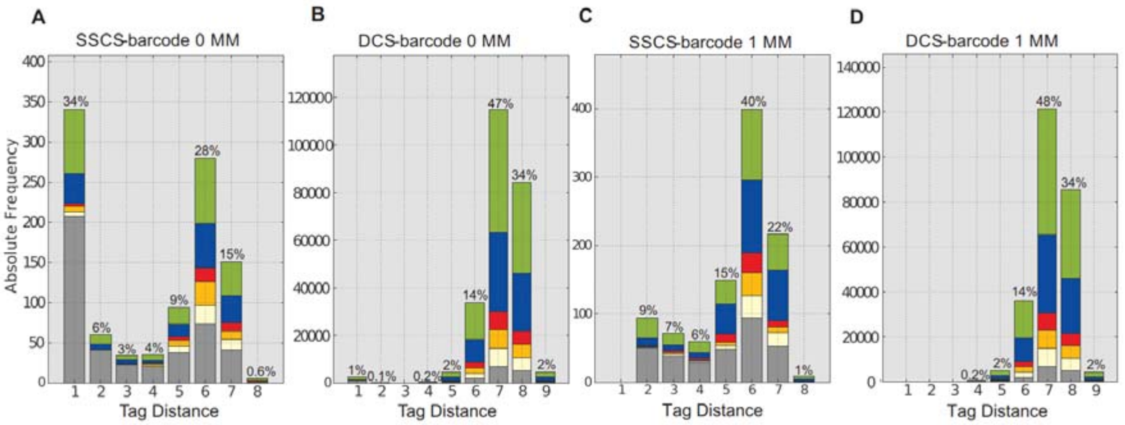

**Supplemental Figure S3.** Tag distance (TD) estimations of the PF2-Standard library. Data are shown before barcode correction. **A.** SSCS tag distance distribution and **B.** DCS tag distance distribution without barcode correction. **C.** SSCS tags and **D.** DCS tags after barcode correction with 1 mismatch. As expected, tags with TD=1 were re-assigned and therefore, the distribution of the TD shifted such that most of the tags differed between 5-6 nucleotides.

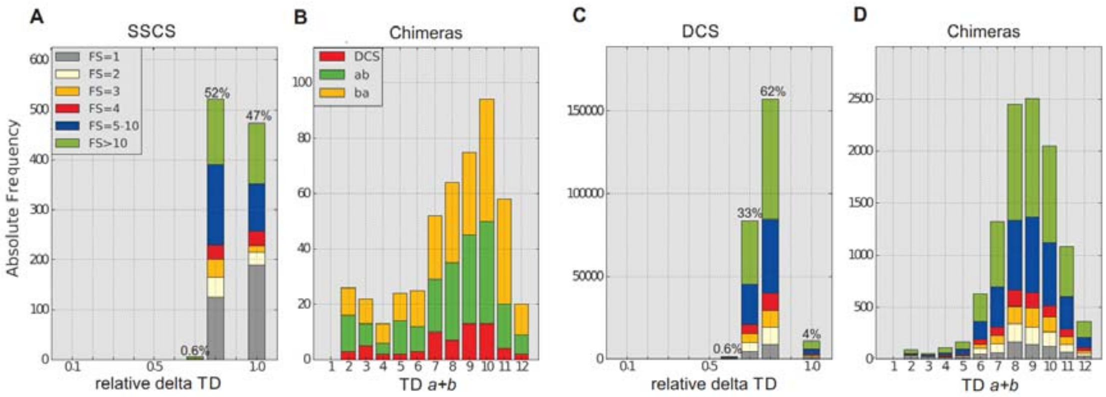

**Supplemental Figure S4. A.** Chimera analysis (CA) of a subset of tags (n=1000) from SSCS the PF2-Standard library after barcode correction (allowing one mismatch). Chimeras have a relative delta TD = 1. **B.** Distribution of TD in chimeric tags (with a relative delta TD of one) stratified into DCS or SSCS. Out of 473 chimeric reads, 185 (39%) belong to SSCS *ab*; 224 (47%) to SSCS *ba*, and 648 (14%) to DCS. **C.** Chimera analysis using the tags of DCS and **D.** TDs of the chimeric DCS.

### Supplemental Notes

#### Supplemental Note 1:

Guidelines for using the QC-tools in Du Novo (Stoler et al. 2016). The full workflow is depicted in Supplemental Fig S1 and can be found under the link https://usegalaxy.org/u/jku-itb-lab/w/paperworkflow-du-novo-20.

##### TD: Tag distance analysis of duplex tags

Each family represents different molecules and therefore, the tags should differ with multiple nucleotides between each other. The tool *"TD: Tag distance analysis of duplex tags"* (TD) calculates the number of nucleotide differences among the tags, also known as Hamming distance (Hamming 1950). This tool processes tabular formatted files with information on family size, the tag sequence, the labelled strand (*ab* or *ba*) arranged in columns. The input file can be produced in Du Novo by the *"Du Novo: Make families"* tool (Stoler et al. 2016) followed by a series of commands to calculate the family sizes as outlined below.
The TD-tool outputs the minimum tag distance (smallest number of differences) for each tag, either as a histogram categorized after the family sizes or as a family size distribution separated by the tag distances. This is very valuable information when implementing the barcode correction tool and deciding how many mismatches should be allowed. We also allow the user additional filters such as minimum/maximum family size or tags forming a DCS.

*→Run "Du Novo: Make families"*
After obtaining the raw reads in the FASTQ format from the sequencing machine, the reads need to be grouped by their barcode into a family. In Du Novo this step is performed by the tool *"Du Novo: Make families"*. Subsequently, the *"Cut"* tool can be used to extract the first two columns (tag, labelled strand *ab*/*ba*). This file is then sorted by the tool *"Sort"* and used as an input for the tool *"Unique lines"*, which adds an additional column containing the family sizes of the tags.

*→ get Tag Distance (TD) to define barcode correction parameter*
The tool *"TD: Tag distance analysis of duplex tags"* (TD) calculates the number of nucleotide differences among the tags extracted from the paired-end reads (PE-reads). Then in a randomly selected subset of 1000 tags, the difference between tags is estimated by comparing each tag with the tags of the complete dataset. Each tag will differ by a certain number of nucleotides with the other tags; yet this tool uses the smallest difference observed with any other tag. The output of the tool plots the minimum tag distance (smallest number of differences) as a histogram categorized after the family sizes (Fig S5 upper panel) or a distribution of family sizes categorized after the TD (Fig S5 lower panel).

*→ Run "Du Novo: Correct barcodes" and verify it with the TD-tool*
Barcode correction allows to reassign tags with up to 3 mismatches to their "true" families. The tool "*Du Novo: Correct barcodes*" accepts values of 1, 2 or 3 as input, which represents the number of differences or mismatches allowed in the tags. If "1" is selected, then tags with one difference or without a difference are united into the same family. This tag correction can be verified by the same TD-tool (e.g. use output from "*Du Novo: Correct barcodes*" and apply the same processing steps as without barcode correction). Re-running the TD analysis with the corrected tags will show if tag

correction worked properly since all the TDs smaller or equal to the selected parameter in barcode correction will be reduced to zero.
As a guideline: if the average number of differences among tags is between 5-6 nucleotides (usually resulting by tagging libraries with a 10+10 or larger random barcode), then 1 mismatch is a conservative setting in the barcode correction tool. However, if smaller barcodes are used (e.g. 8+8 or 6+6 random barcodes) then the average differences will get smaller and start overlapping with differences introduced by sequencing or PCR mistakes.

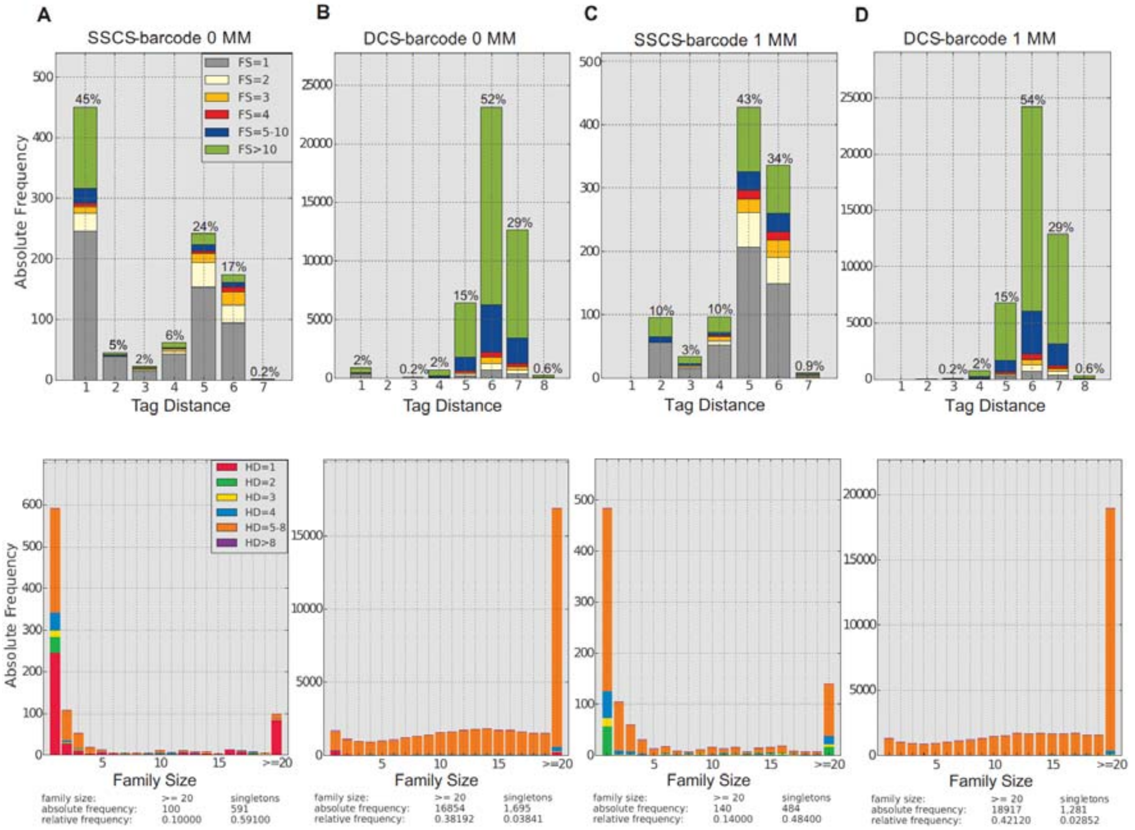

**Supplemental Figure S5. TD distribution of tags** forming **A.** SSCSs or **B.** DCS before implementing the tool "Correct Barcodes" and tags forming **C.** SSCS or **D.** DCS after implementing the tool *"Du Novo: Correct Barcodes"* of PF2-CRISPR library. Barcode correction was performed with one mismatch. The upper panel shows the TD categorized after family size and the lower panel shows the family size distribution (FSD) categorized after TD.

#### Family size distribution (FSD): Family Size Distribution of duplex sequencing tags

The FSD provides a computationally very fast insight into the distribution of the family sizes of ALL tags and a first assessment about the frequency of DCSs at the beginning of the pipeline. This is quite useful in early decision steps of parameters early in the pipeline, such as the minimum number of family members to build the single stranded consensus (SSCS) or the trimming parameters to assure high quality reads. Moreover, this tool can compare several datasets or different steps in the analysis pipeline, such as the effect of tag error correction (e.g. families that were re-united) or loss of families at different steps of the pipeline. In an extension of this tool, each family is stratified into SSCS (*ab*/*ba*) and DSC. This is quite handy to better understand the relationship of SSCS to DCS per family and identify sources of bias (e.g. more SSCS to DCS in a particular family size, or more forward *ab* than reverse *ba* reads). The input file for the FSD-tool is the same as in the TD-tool which contains the family size, labelled strand (*ab, ba*) and the tag.

→ *Run FSD (TD) to assess parameter of family size for SSCS consensus building*

The FSD analyzes the family size associated with a tag, that is the number of reads per tag. It presents this information graphically in several histograms by the tool *"FSD: Family Size Distribution of duplex sequencing tags"* and as a data file (pdf and tabular format). The first part of the output from the FSD-tool "FSD after tags" and "FSD after PE reads" allows the comparison of multiple datasets (e.g. tags before and after barcode correction), from which the user can derive the optimal family size for the alignment of the consensus sequences. Usually, a minimum of 3 reads per family is the requirement in the consensus building and families with only one read (=singletons) or two reads potentially containing valuable information are discarded. The family size distribution helps to decide on the optimal family size parameter for building the SSCS. The same information is also represented based on the PE reads which shows if the data consists of very large families. This means that not only singletons make up a large amount of tags but also large read families (FS>20).

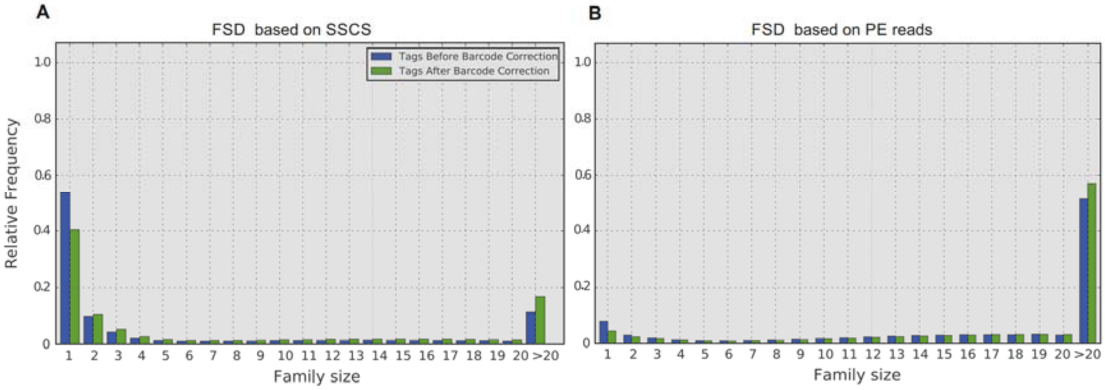

**Supplemental Figure S6. Family size distribution (FSD)** before and after implementing the tool *"Du Novo: Correct Barcodes"* of PF2-CRISPR library based **A.** on tags and **B.** on PE reads. A summary of this figure can be found in Table S5.

In Fig S6 the PF2-CRISPR library before and after barcode correction with one mismatch are compared. Mismatch correction reduces the number of singletons from 54% to 41% which in turn invokes that the number of tags in larger families increased (12% to 17%). But singletons make up only a very small part of the PE reads (8% and 4%, respectively) and halve of the reads are grouped to very large families (FS > 20, 51% and 57%, respectively).

The second part of the output shows the family size distribution (FSD) for each of the datasets in more detail. We categorize the tags based on whether they form DCS (duplex = complementary tags in the *ab* and *ba* direction) or form only SSCS and have no matching complement (only ab or ba direction). The single SSCS-ab and SSCS-ba tags show if a bias between the forward and reverse reads exists. After barcode correction the amount of DCS increases but also the amount of very large families (FS > 20).

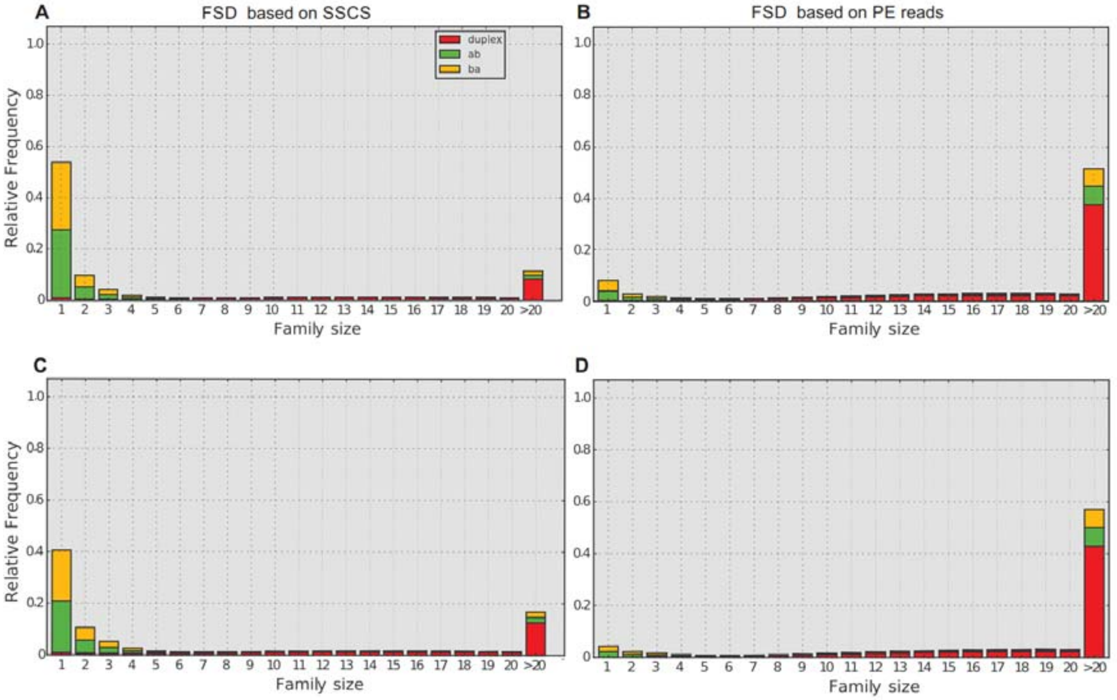

**Supplemental Figure S7. Family size distribution (FSD) A.** before (based on SSCS), **B.** before (based on PE reads) and **C.** after implementing the tool *"Du Novo: Correct Barcodes"* (based on SSCS) and **D.** based on PE reads of PF2-CRISPR library categorized after single SSCS (no matching *ab+ba* tags) and DCS (matching *ab+ba* tags). A summary of this figure can be found in Table S5.

| | #singletons | | | | #FS>20 | | | | | | | |
|---|---|---|---|---|---|---|---|---|---|---|---|---|
| **SSCS before DCS building** | **in SSCS** | | **in PE reads** | | **in SSCS** | | **in PE reads** | | **# tags** | **#PE reads** | **mean FS** | **median FS** |
| before barcode correction | 101,356 | 54% | 101,356 | 8% | 23,306 | 12% | 661,025 | 51% | 188,354 | 1,284,504 | 6.8 | 1 |
| after barcode correction | 56,222 | 41% | 56,222 | 4% | 23,306 | 17% | 733,372 | 57% | 138,631 | 1,284,504 | 9.3 | 2 |

| **single SSCS after DCS building (either SSCS-*ab* or SSCS-*ba*)** | ***ab*** | | | | ***ba*** | | | |
|---|---|---|---|---|---|---|---|---|
| | **in SSCS** | | **in PE reads** | | **in SSCS** | | **in PE reads** | |
| before barcode correction | 72,465 | 38% | 241,042 | 19% | 71,759 | 38% | 238,599 | 19% |
| after barcode correction | 47,132 | 34% | 209,827 | 16% | 46,587 | 34% | 207,247 | 16% |

| **DCS** | **in SSCS** | | **in PE reads** | | **DCS (FS = 1-2)** | | **DCS (FS=3-20)** | | **DCS (FS>20)** | |
|---|---|---|---|---|---|---|---|---|---|---|
| before barcode correction | 44,130 | 23% | 804,863 | 63% | 2,791 | 6% | 25,822 | 59% | 15,517 | 35% |
| after barcode correction | 44,912 | 32% | 867,430 | 68% | 2,312 | 5% | 25,160 | 56% | 17,440 | 39% |

**Supplemental Table S5.** Summarized results of the family size distribution (FSD) (Fig S6 and Fig S7) in SSCSs and PE reads of the PF2-CRISPR library before and after barcode correction.

Fig S7 shows allowing one mismatch in the tag increases the fraction of tags that are able to form a DCS from 23% to 32%. In general, the forward and reverse reads are amplified very evenly during PCR because the number of SSCSs in both directions (*ab, b*a) that cannot form a DCS show similar proportions (38% SSCS-*ab* and 38% SSCS-*ba* before barcode correction, 34% SSCS-*ab* and 34% SSCS-*ba* after barcode correction).

*→ Run "FSD regions" to check enrichment success for different target regions*

The FSD-tool is extended to the *"FSD regions: Family size distribution of user-specified regions in the reference genome"* tool which creates a distribution of family sizes of tags that were aligned to the reference genome. Note that tags that overlap different regions of the reference genome are counted twice for each region. This tool is useful to examine differences in coverage among targeted regions.

Fig S8 includes both complementary SSCS-*ab* and SSCS-*ba* that form a DCS; whereas the Table S6 shows only single counts of the tags per region. The input file for this tool is the same as for the TD-tool which contains the family size, labelled strand (*ab, ba*) and the tag. In addition, a BAM file with the aligned reads to the reference gene/genome and an optional BED file with the chromosome, start and end position of the targeted regions needs to be provided.

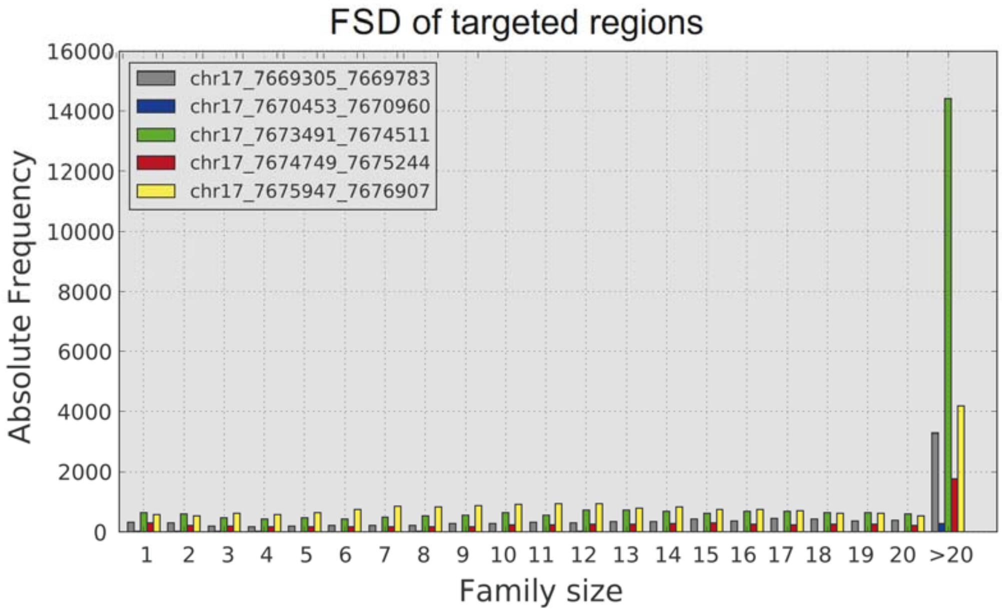

**Supplemental Figure S8. Family size distribution (FSD) distribution grouped by the targeted regions of the PF2-CRISPR library.** All families after barcode correction with FS ≥ 1 that were aligned to the human genome (GRCh38/hg38) are shown (both *ab+ba* tags). Du Novo was performed with a trimming step that allows a minimum read length of 36. Note that all families including the families that overlap to another region are counted. A summary of this figure can be found in Table S6.

| **PF2-CRISPR** | **#SSCS on target** | | **in SSCS-ab** | **in SSCS-ba** |
|---|---|---|---|---|
| chr17:7669305-7669783 | 9,406 | max. FS | 121 | 100 |
| chr17:7670453-7670960 | 480 | #SSCS | 1 (0.04%) | 1 (0.04%) |
| chr17:7673491-7674511 | 26,322 | | | |
| chr17:7674749-7675244 | 6,326 | | | |
| chr17:7675947-7676907 | 19,024 | | | |
| total reads (SSCS) | 1,284,504 | | | |
| total DCS on target regions | 61,558 | | | |

**Supplemental Table S6.** Summarized results of the Family size distribution (FSD) (Fig S8) for the targeted regions of the PF2-CRISPR library after barcode correction with FS ≥ 1. Du Novo was performed with a trimming step that allows a minimum read length of 36. Note that all families including the families that overlap to another region are counted. Therefore, the number of DCSs on the target regions differ to the counts in Table S7. In contrast to Fig S8, only unique tags (either *ab* or *ba* tag of a DCS) were counted. The coordinates are based on the human genome assembly GRCh38/hg38.

*→ Run "FSD Before/After" to check trimming parameters and enrichment success for different target regions*

The tool *"FSD Before/After: Family Size Distribution of duplex sequencing tags during Du Novo analysis"* compares various datasets from the Du Novo analysis Pipeline (Stoler et al. 2016) and helps in decision making of various parameters (e.g. family size, minimum read length, etc.). Du Novo includes a trimming step (*Sequence Content Trimmer*) which can be removed or not. For example: Re-running trimming with different parameters allows to recover reads that would be lost due to stringent filtering by read length. This tool also allows to assess reads on target. The tool extracts the tags of reads and their family sizes before SSCS building, after DCS building, after trimming and finally after the alignment to the reference genome. The first input file contains all unique tags before the consensus building from a TABULAR file which is generated in the same way as in the TD-tool and contains the family size, tag and the labelled strand (*ab/ba*). The second and third input files are the FASTA files from the tools *"Du Novo: Make Consensus Reads"* and *"Sequence Content Trimmer"* (Stoler et al. 2016). And the fourth input file is a BAM file with the aligned reads to a reference genome (e.g. produced by *BWA-MEM* (Li 2013) and *BamLeftAlignIndels* (Garrison and Marth 2012)). In Fig S9, the family sizes for both SSCS-*ab* and SSCS-*ba* are shown; whereas Table S7 represents only counts of either SSCS-*ab* or SSCS-*ba*. In contrast to the *"FSD region"* tool overlapping reads between the regions are counted once and thus, a tag is unique in the data.

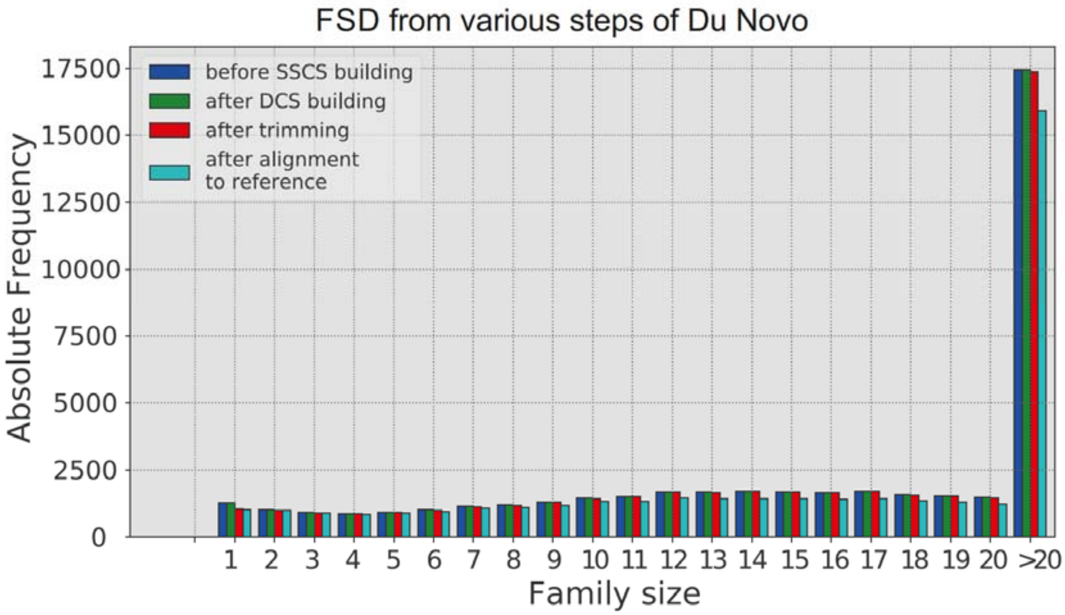

**Supplemental Figure S9.** Family size distribution (FSD) distribution from various steps of the data analysis of the PF2-CRISPR library after barcode correction with FS ≥ 1. Du Novo was performed with a trimming step that allows a minimum read length of 36. Note that in the plot both sizes from the *ab* and *ba* families are represented. As a reference we used the human genome (GRCh38/hg38). A summary of this figure can be found in Table S7.

| PF2-CRISPR | FS ≥ 1 | | in SSCS-ab | in SSCS-ba |
|---|---|---|---|---|
| total reads (all families) | 1284504 | max. FS | 121 | 100 |
| total nr. of families | 116175 | #SSCS | 1 (0.04%) | 1 (0.04%) |
| DCS | 22456 | | | |
| DCS after trimming | 22221 | | | |
| DCS on target | 20049 | | | |

**Supplemental Table S7.** Summarized results of the Family size distribution (FSD) (Fig S9) for some steps of the Du Novo pipeline of the PF2-CRISPR library after barcode correction with FS ≥ 1 Du Novo was performed with a trimming step that allows a minimum read length of 36. In contrast to Fig S9, only unique tags (either *ab* or *ba* tag of a DCS) were counted.

#### Chimera Analysis (CA) in the tool "TD: Tag distance analysis of duplex tags"

→ *check quality of tags by Chimera Analysis (CA)*

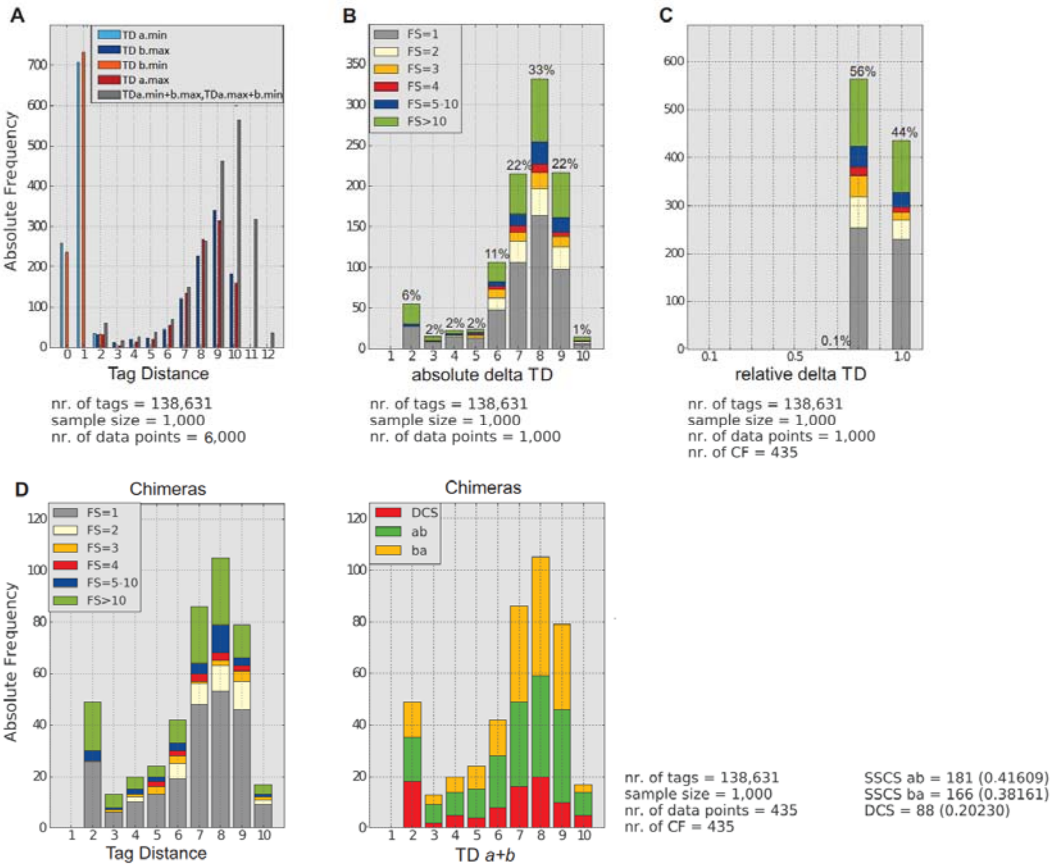

**Supplemental Figure S10.** Summary of the different steps of chimera analysis (CA) after barcode correction of the PF2-CRISPR library. **A.** TD of *a.min*, *b.max*, *b.min* and *a.max* part of the tag (see a detailed description on the calculation in the **Supplemental Note 2**). **B.** Absolute difference between *TD a.min* and *TD b.max* and *TD b.min* and *TD a.max*, respectively. **C.** Relative difference of maximum delta TD. **D.** TD of chimeras categorized by family size or by the formation of DCS (matching *ab*+*ba* tags) or not.

The following steps summarize the *Chimera Analysis (CA)*:

A) Fig S10A shows the differences between the tags at both ends of a read (*a/b*). Chimeras can be distinguished by carrying the same tag at one end combined with multiple different tags at the other end of a read. Here, we describe the calculation of the TDs for only one tag in detail, but the process is repeated for each of the n tags in the random sample (default n=1000) and a detailed description can be found in the **Supplemental Note 2**. First, the tool splits the tag into its upstream and downstream part (named *a* and *b*) and compares part *a* with all other *a* parts of the families in the dataset. Next, the tool estimates the tag distance (TD) among the *a* parts and extracts those tags with the smallest number of differences (TD *a.min*) and calculates the TD of the *b* part. The tags with the largest number of differences are extracted to estimate the maximum TD (TD *b.max*). The process is repeated starting with the *b* part instead and TD *a.max* and TD *b.min* are estimated. Next, we calculate the sum of the TD of both parts.

   Equation 1

   *TD a+b* = TD *a.min* + TD *b.max* and TD *a.max* + TD *b.min*

It is expected that one part of the tag gives much smaller TD (TD = 0-2) than the other because we looked for the minimum TD in one part and for the maximum in the second part (TD = 3-10). Thus, the TD of the whole tag is shifted to larger values (TD = 3-12). We suspected that tags, which are identical in one part of the tag (TD = 0), but different in the second part, are probably artificially introduced.

B) Thus, the absolute difference (= *delta TD*) between the partial TDs are estimated (Fig S10B).

   Equation 2

   *delta TD* = |TD *a.min* - TD *b.max*| *and* |TD *b.min* - TD *a.max*|

   While we expect one of the two halves to be identical in a chimera, the TD of the second half will follow a distribution that depends on the length of the tag sequence. Therefore, the relative difference between the partial TDs, which is estimated in the next step, might be more informative.

C) Fig S10C contains the relative differences of the partial TDs (= *relative delta TD*). Since it is not known whether the absolute difference originates due to a low and a very large TD within a tag or an identical half (TD = 0), the tool estimates the relative TD delta as the ratio of the difference to the sum of the partial TDs. In a chimera, it is expected that only one end of the tag contributes to the TD of the whole tag. In other words, if the same *a* part is observed in combination with several different *b* parts, then one end will have a TD = 0 and therefore not influence the total TD. Thus, the absolute TD difference between the parts (Equation 2) is the same as the sum of the parts (Equation 1*)* or the ratio of the difference to the sum (Equation 3) will equal 1 in chimeric families. Here, the maximum value of the *relative delta TD* and the respective *delta TD* for Fig S10B are selected.

Equation 3

$$relative\ delta\ TD = max\left(\frac{|TDa.min - TDb.\text{max}|}{TDa.min + TDb.max}, \frac{|TDa.max - TDb.\text{min}|}{TDa.max + TDb.min}\right)$$

The plot can be interpreted as follows:

- A low relative difference indicates that the total TD is equally distributed in the two partial TDs. This case would be expected, if all tags originate from different molecules.
- A relative delta TD of 1 means that one part of the tags is identical. Since it is very unlikely that by chance two different tags have a TD of 0, the TDs in the other half are probably artificially introduced and represent chimeric families.

D) Finally, the TD of the chimeras which corresponds to the tags with a relative delta TD of 1 (Fig S10D left). It is also grouped by tags that form a DCS (complementary *ab* and *ba* tags, Fig S10D right). Latter plot is only produced if all tags allowed in the chimera analysis (CA) and only DCS tags.

In our case, ~44% of all tags had a relative difference of 1 which means that one part was identical (TD = 0) to another and the second part was probably artificial introduced (chimeras). About half of the chimeras are singletons that could be removed when a minimum family size of 3 in the consensus building is used.

Chimeras had a rather large TD (TD = 5-10) and we can make sure no sequencing or PCR errors are misclassified as chimeras as we have no chimeras with TD=1. In this sample (n = 1000), around 20% of the chimeras will end up in DCS (red parts) and therefore, might appear as false positives during mutation calling.

In addition to the plots of Fig S10, the chimera analysis produces a tabular file containing the chimeric tags of the sample. Chimeras can be identical in the first or second part of the tag and can have an identical TD with multiple tags. Therefore, the second column of the output file can have multiple tag entries. The file also contains the family sizes and the direction of the read (*ab, ba*). The asterisks mark the identical part of the tag (see an example in Table S8).

| **chimera** | **tags with TD *a.max* or TD *b.max*** |
|---|---|
| GAAAGGGAGG GCGCTTCACG 1 ba | GCAATCGACG ***GCGCTTCACG*** 1 ba |
| CCCTCCCTGA GGTTCGTTAT 1 ba | ***CCCTCCCTGA*** CTCCAATGAC 2 ba,<br>CGTCCTTTTC ***GGTTCGTTAT*** 1 ba,<br>GCACCTCCTT ***GGTTCGTTAT*** 1 ba |
| ATGCTGATCT CGAATGCATA 55 ba, 59 ab | AGGTGCCGCC ***CGAATGCATA*** 27 ba,<br>***ATGCTGATCT*** GAATGTTTAC 1 ba |

**Supplemental Table S8. Example list of chimeras.** The halves of the tags marked with asterisks and in bold show the identical part of the tags. The tags are represented with their family size and direction of the read.

*→ Run Du Novo pipeline + variant calling*

By following this guide the parameters for barcode correction (*Du Novo: Correct barcodes*), consensus building (*Du Novo: Make consensus reads*) and optional trimming (*Sequence Content*

*Trimming*) of the Du Novo pipeline (Stoler et al. 2016) are found. Other steps as the alignment to reference genome (e.g. *BWA-MEM* (Li 2013) and *BamLeftAlignIndels* (Garrison and Marth 2012)) or the variant calling (e.g. *Naive Variant Caller (NVC)* (Blankenberg et al. 2014)) and annotation (e.g. Galaxy tool *Variant Annotator*) are up to the user (see a detailed description of the workflow in Fig S1).

#### Variant Analyzer (VAR-A)

We developed the Variant Analyzer (VAR-A) to assess the evidence supporting a variant call based on a series of different summary data extracted from the raw PE-reads that classifies the confidence level of a variant call by a tier-based system. This allows using more relaxed analysis parameters during consensus building, e.g. small families (including families with only one or two reads) or ad hoc stringent trimming parameters.

VAR-A consists of a series of tools and uses the output of *Variant Annotator, Du Novo: Align families*, as well as the bam files of DCS and SSCS as input.

→ *Run DCS mutations to tags/reads* to extract all tags that carry a mutation in a DCS and create a fastq file of reads with these tags.

→ Trim reads in the fastq file with e.g. Trimmomatic and align the reads to the reference genome e.g. with BWA-MEM.

→ Run *DCS mutations to SSCS stats* to extract all tags from the SSCS bam file that carry a mutation at the same position a mutation is called in a DCS and calculate their frequencies.

→ Run *Call specific mutations in reads* using the files created in the previous steps as input to get the final output.

In the VAR-A output, Each DCS with a variant in the original output is represented by two lines, one for each mate (column "mate"). Information about reads is provided separately for each direction (*ab*/*ba*). The output contains the variant ID, which consists of the position of the variant compared to the reference as well as the reference and alternate alleles, and the tier of the variant is based on the tier classification described in Table 5 and S1. This information is followed by the sequence of the tag (tag), the mate information, the median position within the reads (read pos.ab/ba), the median length of the reads (read median length.ab/ba) and the length of the DCS (DCS median length). Family sizes are reported before (FS.ab/FS.ba) and after QC (FSqc.ab/FSqc.ba), where QC means the removal of reads that could not be aligned to the reference (na.ab/na.ba) or had low PHRED scores at the position of the variant (lowq.ab/lowq./ba). In addition to the absolute number of reads with reference (ref.ab/ref.ba) or alternate (alt.ab/alt.ba), we also report their relative fraction based on the family sizes after QC (rel. ref.ab/rel. ref.ba/rel. alt.ab/rel. alt.ba). The number of SSCS that carry the reference (SSCS ref.ab/SSCS ref.ba) or alternate alleles (SSCS alt.ab/SSCS alt.ba) are also included. If other variants were called within the same family, they are listed in column "other mut". The last column contains possible chimeric tags related to the tag analyzed.

A second tab in the output file provides information about coverage, alternate allele counts (ACs), and allele frequencies (AFs) for each variant based on all tiers (all tiers), high quality tiers (tiers 1.1-2.4), and the original *Variant Annotator* counts (Du Novo). In addition to that, the table contains counts per tier and cumulative allele frequencies using increasing tier numbers.

Finally, in the third tab we supply the total number of variants per tier and a detailed description of the tier based system with examples.

#### Supplemental Note 2:

For a better understanding of this algorithm, we illustrated the process with the following example: For simplicity, suppose a dataset with only five tags, each with a length of 6 nucleotides. The sequence highlighted in grey and yellow represents the *a* and *b* part, respectively:

1. AAAAGTAGGACA (sample tag)
2. ATAACTAGGACT
3. AATACTAGGACA
4. ATTACTAGGACA
5. TTTTCTAGGACA

1. First, we compare the *a* part ($a_1$ = AAAAGT) of the selected tag (AAAAGTAGGACA) with the *a* part of the other four tags.
   The estimated TDs in the *a* parts are TD*a* = 2, 2, 3, 5, respectively (differences labeled in red)
   2. ATAACTAGGACT
   3. AATACTAGGACA
   4. ATTACTAGGACA
   5. TTTTCTAGGACA
2. Next we select the tags with the smallest number of differences (minimum TD) with $a_1$. These are placed in a tag subset called $TDa_{min}$ and include $tag_2$ (ATAACTAGGACT) and $tag_3$ (AATACTAGGACA) with a $TDa_{min}$ = 2.

3. We then compare the b part of the tag ($b_1$ = AGGACA) with the b part of tags in $TDa_{min}$. The b part of $tag_2$ and $tag_3$ differ by distance of 1 and 0, respectively (differences labelled in red). We now select the tag with the largest difference in *b*, which is $tag_2$ with a $TDb_{max}$= 1.
   2. ATAACTAGGACT
   3. AATACTAGGACA

4. We repeat steps 1-3, but this time we start with *b* (TD *b*= 1, 0, 0, 0) and estimate the minimum TD ($TDb_{min}$ = 0, 0, 0) in our dataset ($tag_{3\text{-}5}$)

   2. ATAACTAGGACT
   3. AATACTAGGACA
   4. ATTACTAGGACA
   5. TTTTCTAGGACA

5. Again, we select only the tags that contributed to the minimum TD with $b_1$ ($tag_{3\text{-}5}$) and now calculate the TD for *a* (TD *a*=2, 3, 5) and select the maximum value ($TDa_{max}$= 5).
   3. AATACTAGGACA
   4. ATTACTAGGACA
   5. TTTTCTAGGACA

We ended up with the following TDs from step 1 and 2: $TDa_{min}$=2, $TDb_{max}$= 1
and from step 4 and 5: $TDb_{min}$ = 0, $TDa_{max}$= 5.
This approach presents the basis for the identification of chimeric reads in our dataset, because the TD in chimeric families is expected to be very different between its parts. To assess the difference between parts, we estimate *delta TD:*

$delta\ TD = |TDa_{min} - TDb_{max}|$ and $|TDa_{max} - TDb_{min}|$

In chimeric families the difference (= *delta TD*) between $TD_{min}$ and $TD_{max}$ should be large. Given the large excess of random barcodes to input targets, different molecules should have similar values of $TD_{min}$ and $TD_{max}$. We thus estimate *delta TD* to identify chimeras. We return now to our previous example, and get the following *delta TD*:

*delta TD* =|2 - 1| = 1 and |0 - 5| = 5

At this point, only one comparison renders a large delta TD with a value of 5 ($TDa_{max}$ - $TDb_{min}$); whereas, the other comparison renders small value. In order to normalize these two numbers, we compare *delta TD* with the expected TD of the complete tag (sum of both parts TD *a*+*b*) and estimate the *relative delta TD*. For chimeras, we expect *delta TD* to be equal to TD *a*+*b* (*relative delta TD = 1),* since the TD is contributed only by one of the ends, as the other part is identical (TD = 0).

In our example, we get

$TD\ a+b = TDa_{min} + TDb_{max} = 2 + 1$ and $TDb_{min} + TDa_{max} = 0 + 5$

$$relative\ delta\ TD = max\left(\frac{|TDa.min - TDb.\text{max}|}{TDa.min + TDb.max}, \frac{|TDa.max - TDb.\text{min}|}{TDa.max + TDb.min}\right) = \max\left(\frac{1}{2+1}, \frac{5}{0+5}\right) = 1$$

A relative difference of 1 means that these are chimeric families. Since it is very unlikely that by chance two different tags have a TD of 0 between one of their halves, the TDs in the other half are probably artificially introduced (chimeric reads) and both tags have the same origin even if the TDs is fairly large.

In our example, we have identified the sample $tag_1$ (AAAAGTAGGACA) as a chimeric family. Note that, if we stopped after the first round of calculation for the partial TD ($TDa_{min}$=2, $TDb_{max}$= 1, *relative delta TD*=$\frac{1}{3}$), we would not have identified the tag as chimeric. The *relative delta TD* of 1 was identified in the difference of the TDs from the second comparison ($TDb_{min}$ = 0, $TDa_{max}$= 5, *relative delta TD*=1).
