## Supplementary figures and images for "Increased yields of duplex sequencing data by a series of quality control tools"

### Figure S1

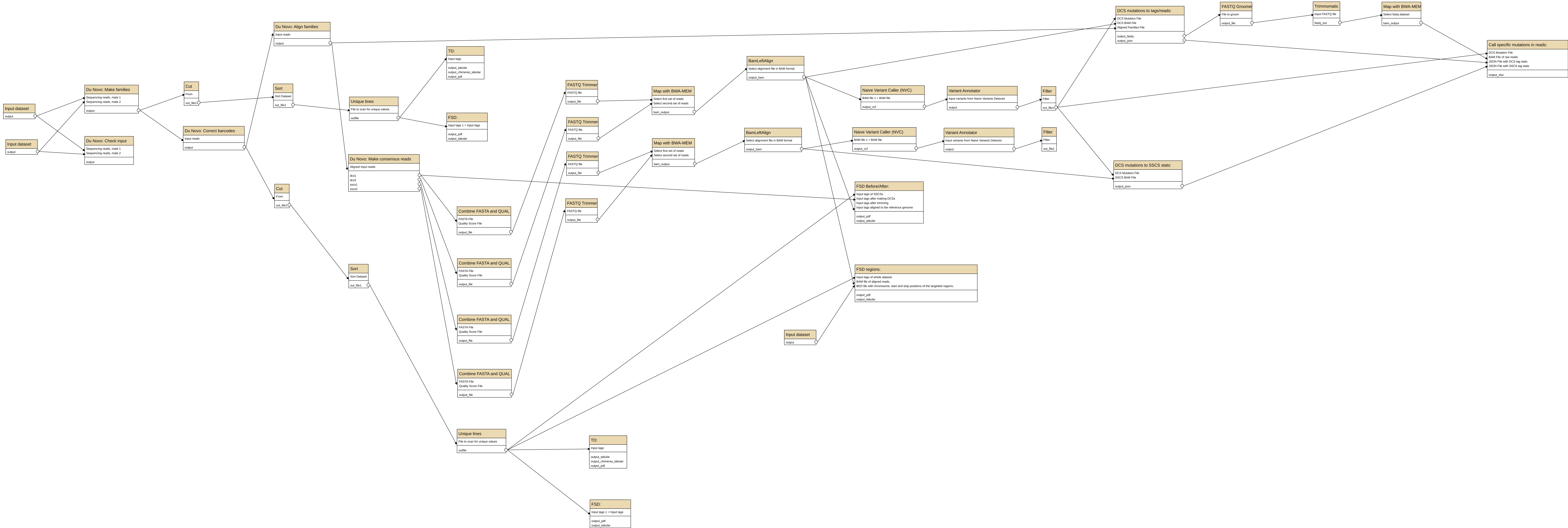
